# Oxytocin Neurons Enable Social Transmission of Maternal Behavior

**DOI:** 10.1101/845495

**Authors:** Ioana Carcea, Naomi López Caraballo, Bianca J. Marlin, Rumi Ooyama, Justin S. Riceberg, Joyce M. Mendoza Navarro, Maya Opendak, Veronica E. Diaz, Luisa Schuster, Maria I. Alvarado Torres, Harper Lethin, Daniel Ramos, Jessica Minder, Sebastian L. Mendoza, Shizu Hidema, Annegret Falkner, Dayu Lin, Adam Mar, Youssef Z. Wadghiri, Katsuhiko Nishimori, Takefumi Kikusui, Kazutaka Mogi, Regina M. Sullivan, Robert C. Froemke

## Abstract

Maternal care is profoundly important for mammalian survival, and non-biological parents can express it after experience with infants. One critical molecular signal for maternal behavior is oxytocin, a hormone centrally released by hypothalamic paraventricular nucleus (PVN). Oxytocin enables plasticity within the auditory cortex, a necessary step for responding to infant vocalizations. To determine how this change occurs during natural experience, we continuously monitored homecage behavior of female virgin mice co-housed for days with an experienced mother and litter, synchronized with recordings from virgin PVN cells, including from oxytocin neurons. Mothers engaged virgins in maternal care by ensuring their nest presence, and demonstrated maternal behavior in self-generated pup retrieval episodes. These social interactions activated virgin PVN and gated behaviorally-relevant cortical plasticity for pup vocalizations. Thus rodents can acquire maternal behavior by social transmission, and our results describe a mechanism for adapting brains of adult caregivers to infant needs via endogenous oxytocin.

**One Sentence Summary:** Mother mice help co-housed virgins become maternal by enacting specific behaviors that activate virgin oxytocin neurons.

## Main Text

Social interactions, such as pair bond formation and child rearing, are fundamental aspects of animal and human behavior (*1–4*). Parental care is especially important in mammals, and new parents must rapidly and reliably express a number of behaviors required for survival of offspring. Some parental behavior is therefore believed to be at least in part innate and hard-wired, or gated by neurochemical changes after mating. However, maternal behavior can also be acquired from experience. In humans and other primates, individuals other than the biological parents can learn to successfully care for children after instruction or observation of experienced caretakers and infants (*1–8*). It is unclear how expression of such alloparenting behaviors in rodents or other species can also be learned from experience, and if so, what mechanisms of neuromodulation and plasticity underlie learning of maternal behaviors.

One of the most important molecular modulators of neural circuit function for social interactions and maternal physiology is the evolutionarily-ancient peptide hormone oxytocin (*1,2, 9,10*). In mammals, oxytocin is released from the hypothalamus and is critical for childbirth and lactation (*10, 11*). Oxytocin also acts in the brain where it is believed to increase the salience of social information, enhancing pair bonding and maternal behavior (*1,9,12-15*), and enabling onset of alloparenting in mice. Specifically, pup-naïve virgin female mice initially ignore neonates and cues related to infant need, e.g., ultrasonic distress calls emitted by pups isolated from the nest (*15*). However, after several days of co-housing with experienced mothers (‘dams’) and litters, most virgin females become maternal and express alloparenting behaviors such as retrieving isolated pups back to the nest. Oxytocin accelerates the onset of pup retrieval in virgin females, promoting plasticity in the virgin auditory cortex to increase the reliability of cortical responses to pup call sounds (*15*). However, little is known about when or how oxytocin neurons are naturally activated to promote pro-social behavior, especially in non-lactating adults.

### An integrated system for monitoring social behavior and neural activity over days

Emergence of pup retrieval in co-housed virgin females provides an opportunity to monitor neural activity including oxytocin neurons during social interactions with adults and infants. We aimed to determine if oxytocin neurons were activated and if behaviorally-relevant cortical plasticity occurred when virgins were co-housed with dams and litters. To examine in high resolution what behavioral events and neural activity patterns led to maternal behavior in virgin female mice, we built a novel integrated system for combined behavioral and neural activity monitoring over prolonged periods (**Fig. 1A**). This system consists of continuous long-term (4 days) videography in the home cage with an overhead camera imaging in both visible (during day) and infrared light (during night), synchronized with audio recordings of vocalizations with ultrasonic microphones, neural recordings from the hypothalamic PVN and auditory cortex, an LED for fiber photometry, and a blue laser for optogenetic identification of optically-tagged oxytocin neurons in vivo.

**Figure 1.**
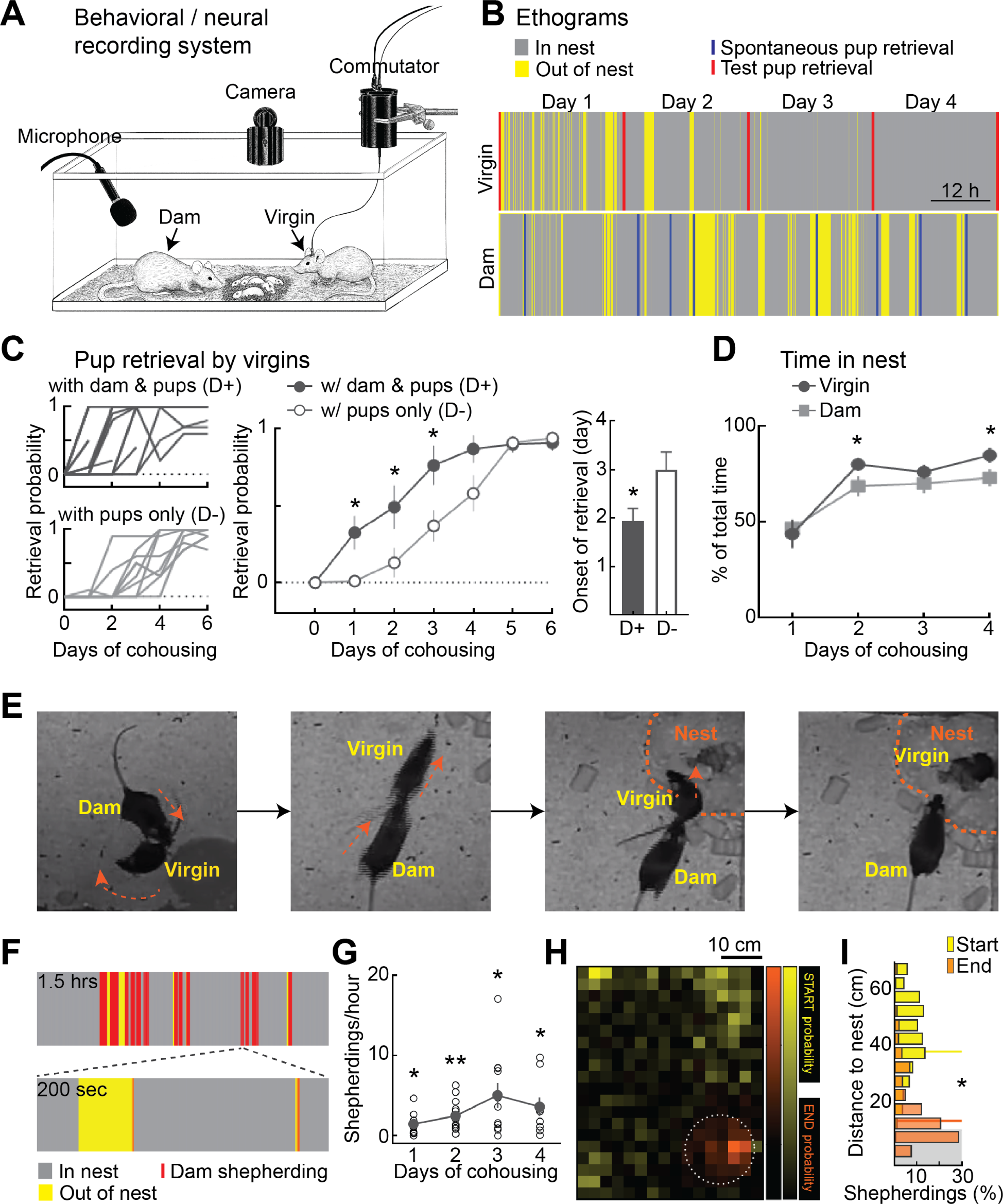
Dams ensure virgins spend time in nest with pups. (**A**) System for multi-day continuous, synchronized monitoring of co-housing behavior, acoustics, and neural activity. (**B**) Ethograms from one virgin-dam pair, including time in nest (gray), out of nest (yellow), spontaneous pup retrieval by dam (blue), and virgin retrieval testing (red). Note change of virgin time in nest on days 3-4. (**C**) Co-housing with dam and pups (‘D+’, N=17 virgins) led to earlier retrieval by virgins than co-housing with pups alone (‘D-‘, N=10 virgins). Left, individual retrieval rates. Middle, average retrieval probability. Right, day of retrieval onset. *, p<0.05. (**D**) Time in nest over days. (**E**) Video frames showing the sequence of shepherding behavior. (**F**) Expanded ethogram showing shepherding events initiating virgin ‘in nest’ residence. (**G**) Shepherding behavior over days. **, p<0.01. (**H**) Probability of shepherding start (yellow) and end (orange) as function of virgin location in home cage relative to nest (dashed circle). (**I**) Distributions of shepherding events start (yellow) and end (orange) distances relative to nest (nest radius was ∼10 cm, gray).

We constructed days-long continuous ethograms (*16*) from the video recordings for each cage of mother, litter, and virgin. We monitored the frequency and duration of specific behaviors of dams and virgins during co-housing (e.g., spontaneous pup retrieval, time spent in nest, **Fig. 1B, Movies S1,S2**). Additionally, we quantified pup retrieval in a separate testing arena at regular intervals over days of co-housing (∼1/day) to determine when virgin females began reliably responding to infant distress calls (**Fig. 1B,C**). We focused on pup retrieval as this form of maternal behavior has several experimental advantages: retrieval is straightforward to score, relies predominantly on a single sensory modality (auditory processing of pup calls), is not initially performed in pup-naïve virgins (0-5% retrieval) but performed essentially perfectly in experienced caregivers (95-100% retrieval), can be expressed rapidly after days of co-housing, and is correlated with the onset of other maternal behaviors such as time spent with pups in the nest (**Fig. 1D**).

We found that the presence of the mother accelerated alloparenting onset in co-housed virgins. Virgins co-housed with an experienced mother and pups began to reliably retrieve on the second day of co-housing, at least a full day earlier than virgins co-housed only with pups (**Fig. 1C, Movies S3,S4**; day one retrieval rate of virgins co-housed with dam and pups: 32.5±10.9%, day one retrieval rate of virgins co-housed only with pups: 1.0±1.0%, p=0.047, ANOVA corrected for multiple comparisons; day two virgins with dam and pups: 49.2±14.0%, day two virgins only with pups: 13.0±9.4%, p=0.025; day three virgins with dam and pups: 76.4±12.6%, day three virgins only with pups: 37.0±10.1%, p=0.013; co-housed with dam and pups: 1.9±0.3 days to retrieval onset, N=17 virgins; co-housed just with pups: 3.0±0.4 days to retrieval onset, N=10 virgins, p=0.03 compared to co-housing with dam and pups, Student’s unpaired two-tailed t-test). This indicated that while all virgins eventually began retrieving, the presence of the mother positively affected retrieval onset in the virgins. Similarly, we noticed that starting on day two, virgins began spending a considerable amount of time in the nest with the pups (**Fig. 1B,D**; dam relative time in nest day one: 46.8±4.6%, day two: 68.8±5.4%, day three: 70.2±5.0%, day four: 73.1±4.5%; virgin relative time in nest day one: 43.8±7.4%, p=0.461 compared to dams; day two: 80.2±3.3%, p=0.031; day three: 76.2±3.7%, p=0.238; day four: 84.9±4.0%, p=0.029; N=13 dam-virgin pairs).

### Mother mice shepherd virgins to the nest for co-caring

What interactions between adult females might lead to earlier expression of maternal behavior in the virgins? We analyzed four days of video per cage and observed two main behaviors performed by the dams. First, we discovered that mothers attempted to keep the virgins within the nest area with the pups. On numerous occasions, if the virgin left the nest, the mother would chase or ‘shepherd’ her back to the nest (**Fig. 1E-G, Movie S5, S6**). This could happen hundreds of times over days of co-housing (**Fig. 1G, fig. S1A**; frequency of shepherding behavior over days, day one: 1.4±0.4 events/hour, p=0.027, Student’s two-tailed t-test; day two: 2.4±0.6 events/hour, p=0.002; day three: 5.0±1.6 events/hour, p=0.012; day four: 3.6±1.2 events/hour, p=0.021; N=13 dam-virgin pairs). We quantified the positions of the virgin when this behavior by the dam would start and end, and found that mothers interacted in this way when the virgin was distal from the nest, and ceased pursuit when the virgin returned to the nest (**Fig. 1H,I**; distances from nest at start of shepherding: 40.5±0.5 cm; distances from center of nest at end of shepherding: 15.8±0.5 cm, p=0.0001, Wilcoxon matched-pairs).

This behavior emerged over hours to days, and persisted over co-housing (**Fig. 1F,G**). The frequency of shepherding events tended to increase (**Fig. 1G**), and the latency from virgin nest exit decreased over the four days of co-housing (**fig. S2**). This seemed quite different from maternal aggression. Indeed, rather than preventing the virgin from approaching pups, the resident dam encouraged the virgin to enter and remain in the nest. Shepherding rarely occurred without pups present (**fig. S1A,B**), so is unlikely to be the dam ‘retrieving’ the virgin to the nest.

### Shepherding and nest entry activate virgin PVN oxytocin neurons

As oxytocin also accelerates pup retrieval onset (*15*), we wondered if interactions with dams or pups naturally activated the hypothalamic oxytocin system in virgins. We recorded from PVN neurons in virgins implanted either with two 8-wire electrode bundles or with 16-channel silicon probes. We recorded from optically-identified oxytocinergic neurons (OT-PVN cells) in Oxt-IRES-Cre mice expressing ChETA channelrhodopsin-2, and in OT-channelrhodopsin-2 mice obtained by cross breeding Oxt-IRES-Cre and Ai32 mice (**Fig. 2A,B**). We confirmed that channelrhodopsin was specifically expressed in OT-PVN cells in transgenic mice (**Fig. 2A, fig. S3**), and that electrodes were implanted in PVN with co-registered micro-computed tomography (μ-CT) and MRI (**Fig. 2A, fig. S4**). We recorded PVN activity in virgins during co-housing and social interactions from 595 PVN single-units in 17 virgins, including 21 optically-identified OT- PVN units, and ensured synchronization of neural, video, and audio recordings (**Fig. 2B**). We also recorded from PVN cells during pup retrieval testing or other interactions with mother or pups to more generally characterize the ‘social receptive fields’ of these units (**fig. S5A,B**).

**Figure 2.**
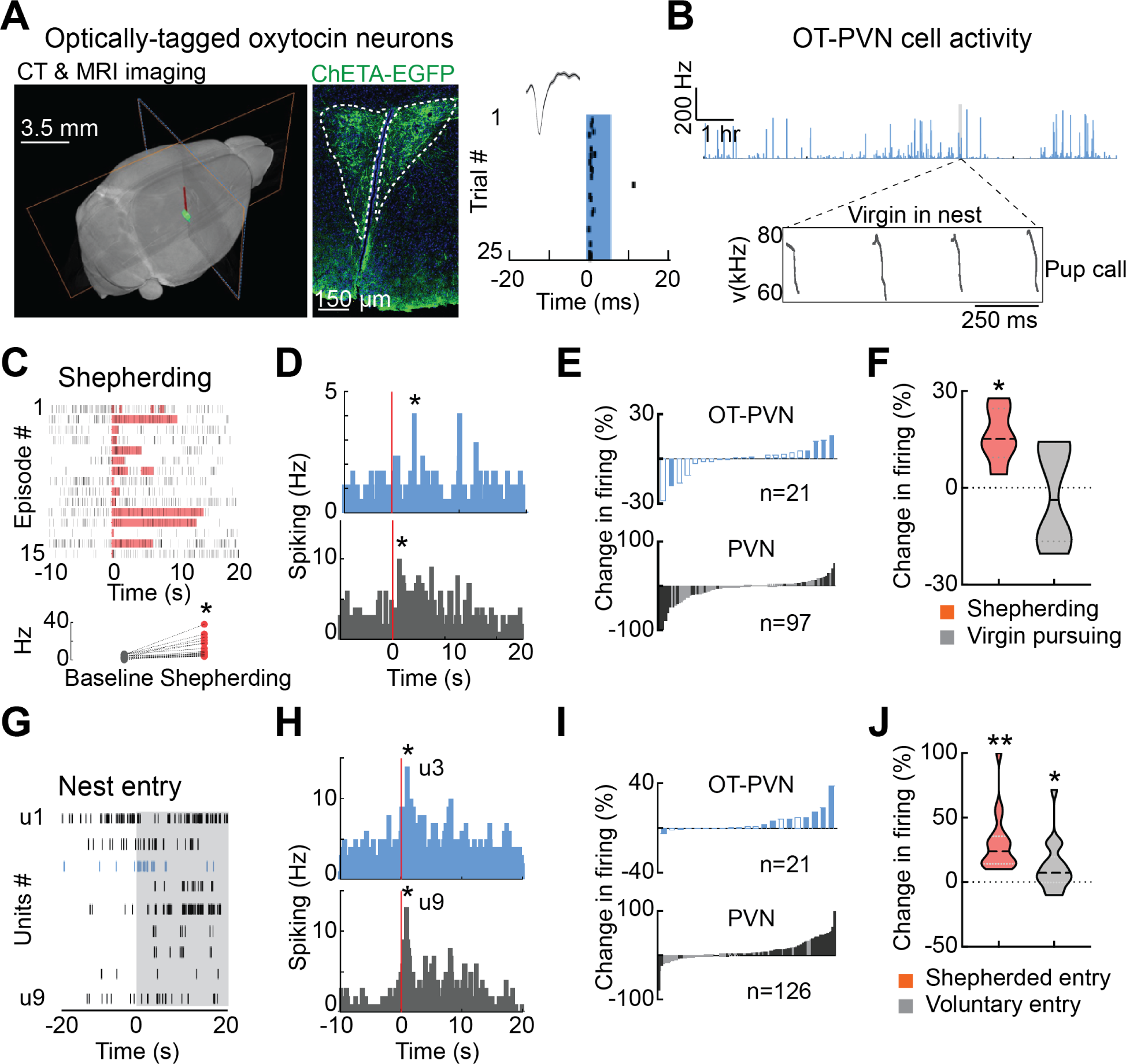
Shepherding and nest entry activated PVN/OT-PVN neurons in co-housed virgins. (**A**) Optically-tagged OT-PVN neuron in vivo. Left, combined MRI/µ-CT for localizing electrodes in PVN. Middle, section from *Oxt-IRES-Cre* virgin female expressing ChETA-EGFP (green) in OT- PVN neurons; DAPI co-stain (blue) for cell number. Right, raster plot of OT-PVN neuron; blue light evoked reliable spikes with short latency. (**B**) OT-PVN neuron recording (top), synchronized with homecage behavior and pup vocalizations (bottom). (**C**) PVN single-unit from co-housed virgin during shepherding. Top, rasters with shepherding periods at time 0 (duration highlighted for each episode). Bottom, increased spiking across episodes for this unit (baseline firing rate: 3.5±0.5 Hz, during shepherding: 12.4±2.2 Hz, p=0.0001). (**D**) Peri-event time histograms of example OT-PVN (top, blue) and PVN (bottom, gray) neuron activity during shepherding starting at time 0; *, significant bursting. (**E**) Change in spike rate during shepherding for individual units (filled bars significantly different during shepherding; open bars, not significantly different). (**F**) Change in spike rate during shepherding (red) was significantly higher than episodes where virgins pursued dams (gray). (**G**) Nine simultaneously-recorded PVN cells including OT-PVN cell (u3, blue) during nest entry event. (**H**) Peri-event time histograms of example OT-PVN (top) and PVN (bottom) neuron activity during nest entry at time 0. (**I**) Change in spike rate during nest entry. (**J**) Increase in spiking when virgin entered the nest after shepherding (red) or voluntarily (gray).

We found that when the mother shepherded the virgin back to the nest, a specific subpopulation of neurons in virgin PVN were transiently activated, including optically-identified OT-PVN neurons (**Fig. 2C-E, fig. S6A**; 4/21 OT-PVN neurons and 12/97 PVN neurons recorded during these interactions significantly increased activity). The increased spiking of these neurons was not due to movement, as we did not observe a similar modulation of activity during episodes where the virgin pursued the mother (**Fig. 2F**; increase during shepherding: 16.1±3.4%, decrease during pursuing dam: –2.4±6.4%, p=0.03), a behavior equivalent in motor but not social aspects. The moment of nest entry was a particularly powerful stimulus that activated PVN and OT-PVN neurons, both when virgins were shepherded into the nest as well as when virgins voluntarily entered the nest (**Fig. 2G-J, fig. S6B**; 7/21 OT-PVN neurons and 41/126 PVN neurons significantly increased activity at nest entry; after shepherding: 29.5±5.2%, p=0.0001; voluntary entry: 3.4±2.3%, p=0.02, n=18 units recorded in both conditions). However, pups had to be in the nest in order for these cells to fire, as PVN neurons were not activated by entering an empty nest without pups (**fig. S5B**). During these times, the abandoned pups in the nest would emit distress calls (**Fig. 2B, fig. S7**). This might lead to a natural pairing of pup distress calls with PVN activation and oxytocin release in virgin brain areas, including those regions important for processing vocalizations such as the auditory cortex.

### Virgins observed mothers demonstrating maternal behavior

We found that dams would spontaneously retrieve pups during co-housing. During these episodes, dams would drop or drag pups out of the nest and then retrieve them back into the nest (**Fig. 3A, Movie S1,S7**). This occurred on average once every two hours throughout co-housing per mother (**fig. S8**), and co-housed virgins would be able to observe these self-generated maternal retrievals. Remarkably, these retrieval episodes performed by the mothers evoked responses in virgin PVN (**Fig. 3B,C, figS9**; 5/17 OT-PVN neurons and 12/69 PVN neurons recorded in virgins during these episodes significantly increased activity when dams retrieved pups), even though the virgin was not directly interacting with dam or pups. More rarely, we found that the mother would deposit pups in front of the virgin (**Movie S8**). Even if the virgin did not retrieve the pups brought to her, virgin PVN neurons could increase firing during these episodes (**fig. S10**). These interactions would also lead to a natural pairing between oxytocin activity and pup call sounds made by the isolated pups directly in front of the virgin.

**Figure 3.**
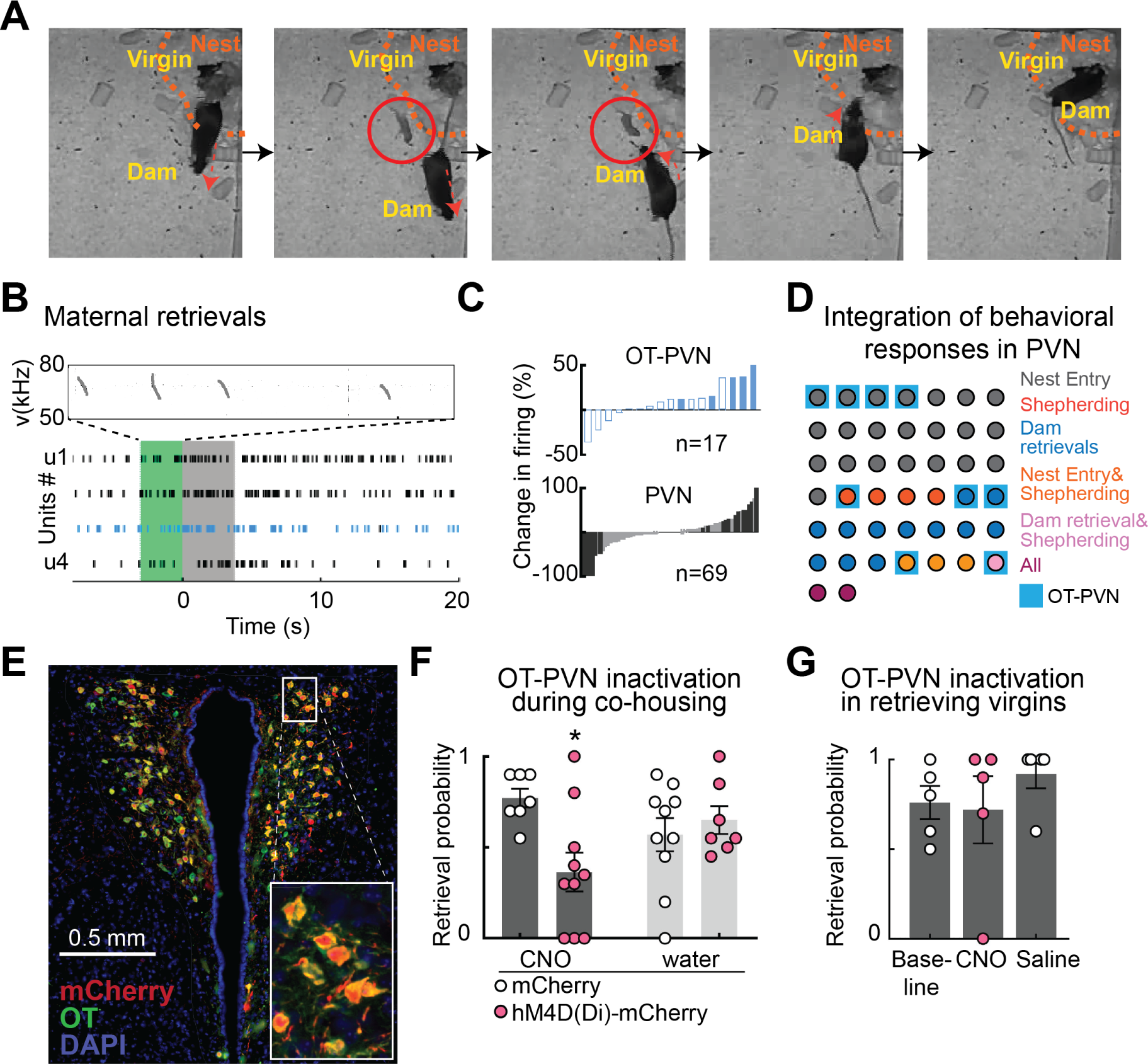
Spontaneous pup retrieval by dams. (**A**) Video frames of dam moving pup out of nest and retrieving it back to nest in front of virgin. (**B**) Four simultaneously-recorded virgin PVN cells including OT-PVN cell (blue) during spontaneous retrieval episode by dam. Green, pup isolation after dropping by dam (inset, pup isolation calls); gray, dam retrieves pup. (**C**) Change in spike rate in virgins during spontaneous retrievals by dams. (**D**) Heterogeneous responses of those PVN units that were monitored during nest entry, shepherding, and maternal retrieval, and had significant responses during at least one of those behaviors. Identified OT-PVN units that met these criteria are highlighted in blue. (**E**) DREADDi-mCherry expressed in OT-PVN cells in *Oxt- IRES-Cre* mice. (**F**) CNO treatment in DREADDi-expressing mice during initial co-housing reduced retrieval. Viral expression alone, without CNO, did not significantly affect retrieval. (**G**) CNO in retrieving females that express DREADDi did not affect retrieval.

Thus we identified three specific behaviors encouraged or engaged by the dams that activated virgin PVN neurons: shepherding, nest entry, and maternal retrievals. Do these behaviors engage same or different neuronal populations? A total of 44 units (including 9 OT-PVN units) showed significant responses to at least one behavior and were also recorded during displays of all three types of behavior (**Fig. 3D**). A substantial proportion of PVN neurons were activated exclusively by nest entry, whereas smaller subsets were activated exclusively by shepherding or by dam retrievals. A smaller number of units were activated by two or all three behaviors, but overall the population of PVN neurons displayed considerable heterogeneity in terms of response selectivity.

If PVN oxytocin neuron firing is a mechanism for accelerating pup retrieval in the co-housed virgins, then inactivating these cells should slow or prevent retrieval onset. To silence these cells during co-housing, we expressed inhibitory DREADD receptors in *Oxy-IRES-Cre* mice with AAV2/2-Syn:DIO-hM4D(Di)-mCherry (**Fig. 3E**). Administering the DREADD ligand CNO to specifically reduce firing of oxytocin neurons led to delayed retrieval onset in DREADDi- expressing naïve virgins compared to animals receiving water or virgins injected with a control virus (AAV2-Syn:mCherry) receiving the same CNO treatment (**Fig. 3F**; ‘mCherry+CNO’ retrieval after first two days of co-housing: 77.1±5.1%, N=7; ‘hM4D(Di)-mCherry+CNO’: 36.5±10.6%, N=10; ‘mCherry+water’: 57.9±9.1%, N=10; ‘hM4D(Di)-mCherry+water’: 65.0±7.6%, N=7; two-way ANOVA, treatment F(1,30)=0.2, virus F(1,30)=3.1, interaction F(1,30)=6.8, p=0.0135). However, after virgins began retrieving, CNO administration did not significantly affect pup retrieval (**Fig. 3G**; baseline retrieval: 76.0±9.2%, N=5; CNO: 72.0±18.8%, N=5; saline: 92.0±8.0%, N=5; repeated measures ANOVA, p=0.332). This shows that activity of hypothalamic oxytocin neuron firing is required during the first few days of co-housing to initiate alloparenting behavior, but that the oxytocin system is no longer required to maintain retrieval after it is learned.

### Social transmission of maternal behavior by observation

We hypothesized that the onset of retrieval behavior in virgins was correlated with exposure to the number of retrievals performed by the dam. If so, then explicitly testing the dam in the presence of the virgin might also accelerate virgin retrieval onset. In a separate cohort of non-co-housed virgins, we first performed daily 10 retrieval tests with the dam for four days, with the virgin present in the arena during the retrieval testing. Retrieval abilities were then also assessed daily in each virgin 30 minutes later without the dam present (**Fig. 4A**). We observed that the onset of retrieval behavior in the virgins increased as a function of the total number of pup retrievals performed by the dam in the presence of the virgin (**Fig. 4B**, ‘No barrier’; virgins retrieving after 40 maternal demonstrations: 11/15 virgins retrieving).

**Figure 4.**
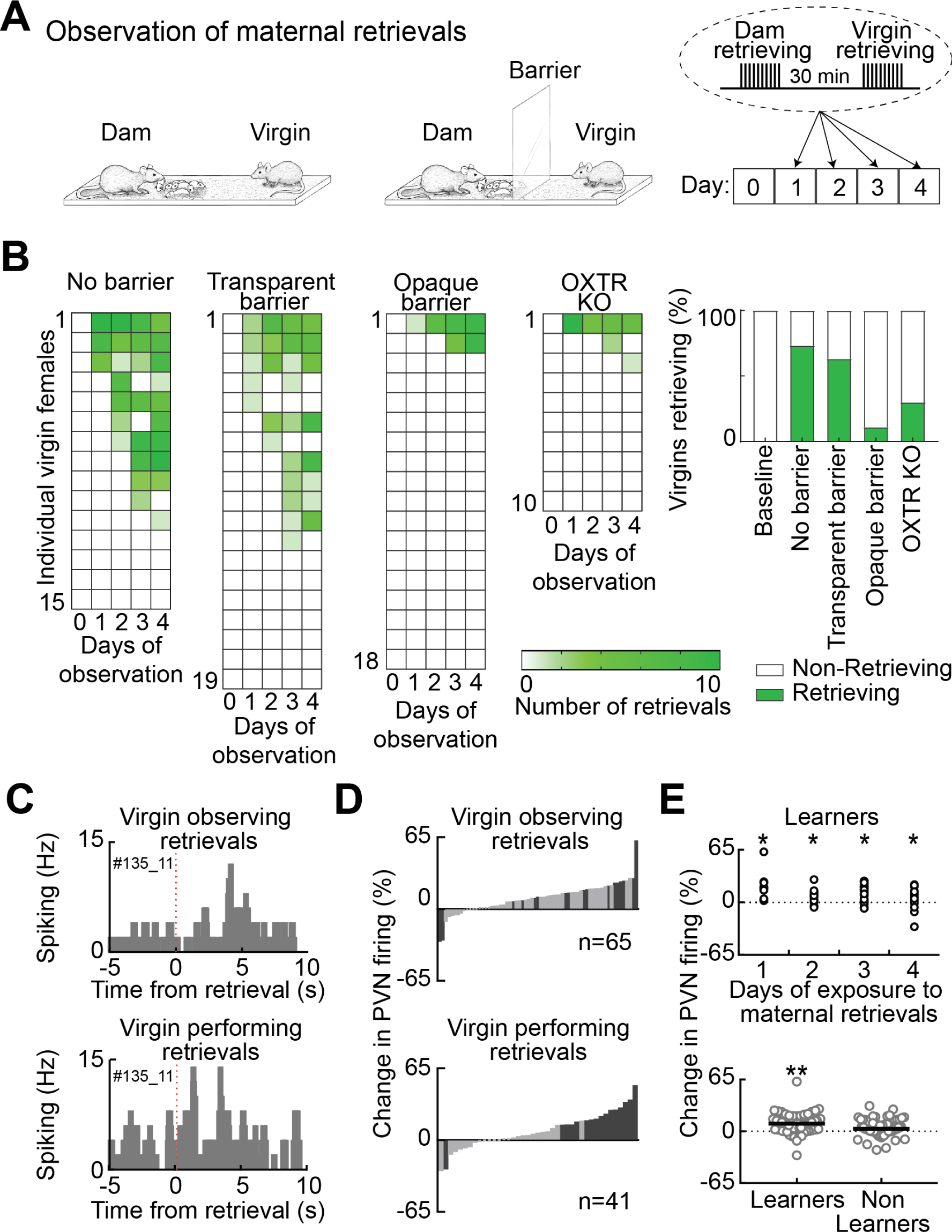
Observational learning of pup retrieval. (**A**) Schematics of non-co-housed retrieval testing; some cages contained plexiglass barriers. (**B**) Non-co-housed virgin retrieval increased with observation when virgins were in cage during dam retrieval without (left)/with transparent barrier (center). Virgins did not begin retrieving when barrier was opaque. Observational learning required oxytocin signaling, as oxytocin receptor knockout mice did not begin retrieving. (**C**) Virgin PVN unit responding to both observation of maternal retrieval (top) and self-performance of pup retrieval (bottom); spike trains aligned to retrieval start at time 0. (**D**) Change in spike rate in virgins during observation of retrievals by dams (top; filled bars, units that significantly changed firing) and during execution of retrieval (bottom). (**E**) PVN activity during observation predicted learning. Spiking significantly increased during observed maternal retrievals on all days in learners (top), but not in animals that did not learn to retrieve (bottom).

We wondered if this depended on physical contact, and so we tested a separate group of pup-naïve virgins in split cages with a barrier between the virgin and where the dam was tested for pup retrieval. However, most virgins behind a transparent barrier also learned to retrieve (**Fig. 4B**, ‘Transparent’, **Movie S9**; retrieval after 40 demonstrations: 12/19 virgins retrieving). To test if visual observation of the retrieving dam was required, we made the barrier opaque in a third group of naïve virgins. In striking contrast to the transparent barrier, virgins did not begin retrieving pups at all when prevented from directly observing retrieving dams (**Fig. 4B**, ‘Opaque’, **Movie S10**; retrieval after 40 demonstrations: 2/18 virgins retrieving). This form of socially-transmitted maternal behavior also required oxytocin signaling. Retrieval in virgin oxytocin receptor knockout mice was negligible over all four days and was not enhanced by observing maternal retrievals (**Fig. 4B**, ‘OXTR-KO’; retrieval after 40 demonstrations: 3/10 virgins retrieving). Thus, more virgins learned to retrieve pups after observing dam retrievals unhindered or through a transparent barrier, compared to when obstructed by an opaque barrier or when missing OXTR (**Fig. 4B**, right: Chi-square, p<0.0001). As rates of retrieval learning in non-co-housed virgins are lower than those occurring during a few days of cohousing (**Fig. 1C**), it is likely that each of the different behaviors enacted by the dam and virgin may help promote successful maternal care by the co-housed virgins. This might be due to the heterogeneity of the oxytocin system (**Fig. 3D**), in terms of which behaviors or experiences activate which subset of cells at a given time.

These data indicate that virgins can learn alloparenting abilities from observing experienced females. The behavior of the isolated pup during observation did not greatly contribute to how well virgins learned to retrieve, as pup movement and vocalizations were similar between virgins who ended up learning and those who did not learn to retrieve (**fig. S11**). If indeed pup retrieval is at least partially observationally learned from experience, virgins might first make errors in trying to retrieve pups before successfully performing each of the components of retrieval: searching for the pup based on acoustic cues, rotating the pup to gain access to the back of the pup, lifting the pup from the ground without causing pain, and returning the pup to the nest. We examined movies of virgin females that began retrieving, and found that before their first successful retrieval, some animals made errors (including examining or sniffing pups without retrieval, aborted retrievals, and improper motor actions such as biting head or tail) consistent with trial-and-error learning (**fig. S12, Movie S9**).

Retrievals by the dam activated virgin PVN neurons, even though the virgins themselves were not otherwise interacting with the dam or pup (**Fig. 4C-E**). When virgins had learned to retrieve, we found that several of the same cells active during observation also fired during subsequent performance of pup retrieval (**Fig. 4C,D**; top, 15/65 virgin PVN neurons significantly increased activity when dams retrieved pups; bottom, 15/41 of the same PVN units significantly increased activity when virgins retrieved themselves). This elevated activity in PVN predicted which animals would learn to retrieve by observation. Increased PVN activity was highest on the first day of observation (**Fig. 4E**, top), and significantly elevated overall only in those virgins that eventually began retrieving (**Fig. 4E**, bottom, ‘Learners’, activity increase of 9.8±1.5%, N=3 virgins, n=65 units). In virgins who failed to begin retrieving, we did not observe significantly elevated PVN activity (**Fig. 4E**, bottom, ‘Non-learners’, activity increase of 3.6±0.9%, N=5 virgins, n=103, p=0.0001 compared to learners, Mann-Whitney). Thus, there is a shared hypothalamic cell population sensitive to both watching and performing retrieval. This indicates that both forms of learning (during co-housing and via observation) share mechanisms and neural activity patterns for initiating maternal behavior (*17, 18*)

### PVN activity predicts single-trial modulation and plasticity of auditory cortex

These results show that, specific actions of mother mice can accelerate the emergence of maternal behavior in virgins that are exposed to the behavior of the mother. Neurons in the virgin brain are sensitive to these behaviors, leading immediately to PVN activation and consequent oxytocin release throughout the brain, which seems to accelerate alloparenting onset over a longer time period. What changes connect PVN firing and oxytocin modulation to pup retrieval behavior? One important neural adaptation with motherhood occurs within left auditory cortex, which becomes more responsive to pup distress calls and is required for successful retrieval. This plasticity is accelerated by oxytocin signaling within left auditory cortex (*15*). To examine if this plasticity naturally occurs with PVN activation during co-housing behaviors, we recorded from left auditory cortex and PVN of co-housed virgins, with four different methodological approaches. In one group of virgins, tetrodes were implanted in left auditory cortex (**Fig. 5A,C**), in a second cohort fiber photometry was used to assess cortical activity (**Fig. 5B,C**), in a third group of virgins we simultaneously recorded in PVN and performed photometry in cortex (**Fig. 5D,E**), and finally we recorded from virgin PVN neurons projecting to left auditory cortex (**Fig. 5F,G**).

**Figure 5.**
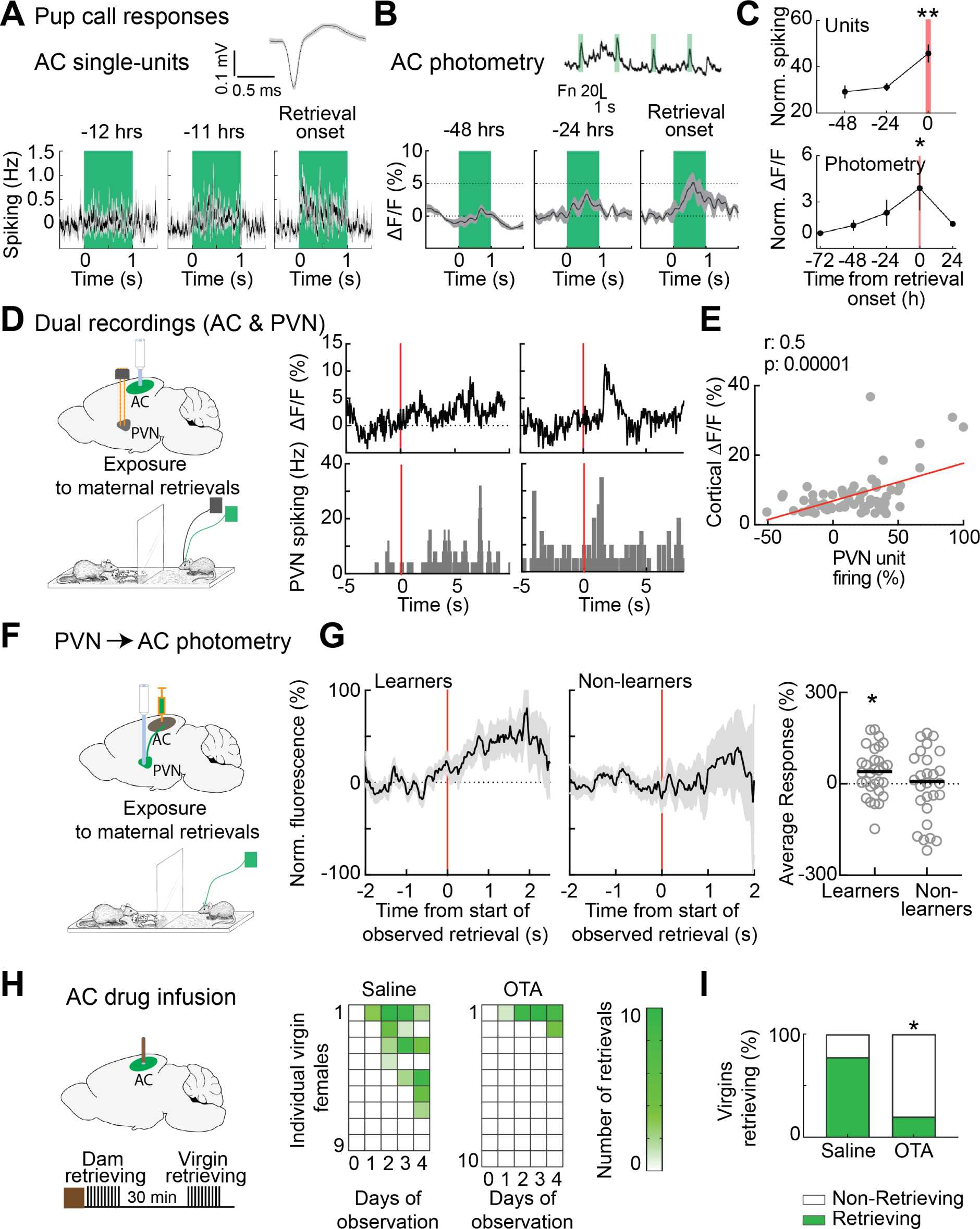
PVN activity predicts plasticity for pup calls in left auditory cortex. (**A**) Single-unit recordings throughout co-housing with tetrodes in virgin female left auditory cortex. Top, example waveform recorded in left auditory cortex. Bottom, example animal which began retrieving 12 hours into co-housing. Note increase in single-unit spiking to pup call (green) prior to retrieval onset. (**B**) Fiber photometry from virgin female left auditory cortex over days of co-housing. Top, example trace from virgin on day of first retrieval. Bottom, example animal which began retrieving 48 hours after start of co-housing. Note increase in photometric response prior to retrieval onset. (**C**) Top, Summary of single-unit recordings with tetrodes in left auditory cortex during co-housing. Bottom, summary of fiber photometry. (**D**) Left, schematic of simultaneous cortical photometry and single-unit recording in PVN during observation of maternal retrievals. Right, example traces from one animal before (left) and after learning to retrieve (right). (**E**) Summary of single-trial modulatory events correlating PVN activity with cortical photometry responses. (**F**) Schematic of photometry recordings in virgins from PVN neurons projecting to left auditory cortex. (**G**) Left, photometry traces aligned to start of observed maternal retrievals in learners (left) and non-learners (middle). Summary of responses averaged over two seconds after retrieval start show significant responses in learners but not non-learners. (**H**) Schematic of ‘Saline’ or ‘OTA’ infusions in the left auditory cortex right before observation of retrieval. OTA but not saline disrupted acquisition of pup retrieval behavior by observation. (**I**) Summary showing that very few virgins injected with OTA in the auditory cortex learned retrieval.

We found that before co-housing, responses to pup calls were minimal in virgin left auditory cortex. However, prior to the first retrieval episode, auditory cortex began responding to pup call sounds, as first measured with single-unit recording to measure the responses of individual neurons (**Fig. 5A**). Emergence of neural pup call responses in left auditory cortex predicted retrieval onset (**Fig. 5A,C**; spiking response to best pup call 48 hours before retrieval onset: 29.3±2.8% increase relative to baseline, n=71 units from 2 animals; spiking 24 hours before retrieval: 31.3±1.7% increase, n=73 units; spiking on day of first retrieval: 45.8±3.9% increase, n=40 units, p=0.005 compared to spiking 48 hours before retrieval onset).

Similar results were obtained using fiber photometry to measure the overall sensitivity of auditory cortex in wild-type virgin female mice virally expressing GCaMP6s in cortical neurons (**Fig, 5B,C**). With photometry we could assess the changes in pup call responses in the same animals over days, and found a progressive increase in photometric signal that predicted the day of retrieval onset (**Fig. 5C**, bottom; dF/F response to best pup call 48 hours before retrieval onset: 1.5±0.3%; dF/F 24 hours before retrieval: 2.3±0.8%; dF/F on day of first retrieval: 3.9±1.4%, p=0.003 compared to baseline signal 72 hours before retrieval onset; N=3 mice).

We asked if the emergence of cortical responses to pup calls relates to PVN activity. We made simultaneous recordings from virgin PVN and auditory cortex, with electrodes implanted in the hypothalamus and a fiber for photometry from cortex (**Fig. 5D,E**); in some cases, the PVN cells specifically projecting to the auditory cortex expressed GCaMP6s (PVN→AC; **Fig. 5F,G; fig. S13**). We took advantage of the trial-based structure of retrieval testing and observational learning in non-co-housed animals separated by a transparent barrier, and simultaneously recorded cortical and PVN signals in the observing virgins, together with audio recordings of pup call sounds during episodes of pup retrieval by the dams. We found that interactions between the dam and virgin could activate PVN tens to hundreds of milliseconds before increasing excitability in auditory cortex (**Fig. 5D**). Prior to retrieval onset, peaks in PVN activity were correlated with pup call-evoked responses in cortex; conversely, failure of PVN activation was correlated with a negligible cortical signal (**Fig. 5E**). When this PVN activation occurred together with detectable pup calls, several episodes of this natural pup call pairing led to persistent enhancement in cortical responses to pup calls. In this way, the newly-parental auditory cortex becomes much more sensitive to pup distress calls, allowing for rapid behavioral responses to emerge as mice learn to retrieve pups (**fig. S14**). This was evident in the photometry from PVN→AC projections, in which PVN→AC neurons in virgins who began retrieving were activated by start of observed dam retrieval (**Fig. 5G**, ‘Learners’, 38.21±14.4%, p=0.01), but virgins that did not learn after observation showed less activation of the PVN→AC projection (**Fig. 5G**, ‘Non-learners’, 4.05±24.16%, p=0.8).

To determine if the enhanced correlation between PVN and cortical activity indicates a role for cortical oxytocin signaling in the observational leaning of pup retrieval behavior, we infused either saline or a selective oxytocin receptor antagonist (‘OTA’) in the auditory cortex. Wild-type females did not begin retrieving when OTA was infused into left auditory cortex of virgins before the observational learning paradigm (**Fig. 5H**, ‘Saline’, virgins retrieving after 40 demonstrations: 7/9; ‘OTA’, virgins retrieving after 40 demonstrations: 2/10; Fisher’s exact test, p=0.023).

## Discussion

Parenting is one of the most important survival-related behaviors, performed by altricial animals (*2,4,19,20*). In many species, although many parental behaviors are assumed to be innate due to their rapid onset and reliable performance required for adequate child care and survival, it has long been appreciated that most behavior results from both intrinsic and acquired processes (*21, 22*). Here we examined the extent to which maternal behavior can be acquired by social transmission. We constructed an integrated system that combined continuous days-long videography with audio and neural activity monitoring, enabling the recording of documentary movies and subsequent behavioral discovery. We found that experienced mothers engage in at least two behaviors that cause oxytocin neurons to fire in the hypothalamus of a co-housed virgin female, and appear related to learning of maternal behavior via visual observation and perhaps teaching. First, the dam shepherds the virgin towards pups in the nest but not towards an empty nest, to help ensure that pups are protected by at least one of the co-caregivers. Second, virgin mice have numerous opportunities to observe how the dam retrieves pups after being displaced from nest. We showed that virgin mice can acquire robust pup retrieval behavior in part by observing maternal retrievals. This observational learning depends on visual input, but could also use auditory and olfactory inputs, as in other forms of observational learning across species (*23–29*).

It has been difficult to directly connect modifications of neural circuits to changes in behavior (*30–32*), for at least two reasons: first, a suitable behavioral model is required for measuring or manipulating neural circuits over the entire time-course of behavioral changes; and second, changes in neural activity may occur rapidly and transiently, at various times for different animals. Furthermore, behaviorally-relevant plasticity in vivo generally requires neuromodulation (such as activation of the oxytocin system), combined with local circuit activity driven by sensory experience (*31, 33*). Our approach here took advantage of the speed and reliability of maternal behavior onset (and pup retrieval in particular), to record directly for the first time from identified oxytocin neurons in non-lactating animals during social interactions. This allowed us to relate episodes of PVN oxytocin neuron firing to moments of modulation and plasticity within left auditory cortex, required for pup retrieval behavior (*15, 17*). It remains to be determined what other sensory cues from the dam and pups such as vocalizations or olfactory signals (*34–37*) might help instruct or incentivize the virgin leading to experience-dependent plasticity required for successful maternal behavior, or which aspects of parental care are instead hard-wired or innate. This presumably involves changes to other aspects of neural function beyond sensory processing, including motor learning required for correct pup retrieval and other forms of parental care.

## Acknowledgements

We thank G. Buzsaki, E. Glennon, V. Grinevich, B. Hangya, K. Kuchibhotla, M.A. Long, J. Minder, J.K. Schiavo, R. Tremblay, R.W. Tsien, and D. Vallentin, for comments, discussions, and technical assistance; and C.A. Loomis and NYU School of Medicine Histology Core for assistance with anatomical studies. Oxytocin-IRES-Cre mice were obtained from The Jackson Lab. Oxytocin receptor knockout mice were obtained from R.W. Tsien (NYU School of Medicine). S.E. Ross created artwork in **Figs. 1A, 4A, 5D,F**. This work was funded by NICHD (HD088411 to R.C.F. and R.M.S.), NIDCD (DC12557 to R.C.F.), the BRAIN Initiative (NS107616 to R.C.F., A.M., and D.L.), NIMH (K99/R00 MH106744 to I.C.; F32 MH112232 to ., and T32 MH019524 to B.J.M.), the Strategic Program for Brain Sciences from Japan (AMED 16K15698 to S.H. and K.N.), a McKnight Scholarship (R.C.F.), a Pew Scholarship (R.C.F.), a Howard Hughes Medical Institute Faculty Scholarship (R.C.F.); and NARSAD Young Investigator awards (I.C. and M.O.). In vivo imaging was performed under the DART Preclinical Imaging Core partially funded by the NYU Laura and Isaac Perlmutter Cancer Center Support Grant, NIH/NCI P30CA016087. The Center for Advanced Imaging Innovation and Research (CAI2R, www.cai2r.net) at NYU School of Medicine is supported by NIH/NIBIB P41 EB017183.

## Author Contributions

I.C., N.L.C., R.O., J.M.M.N., J.S.R., M.O., B.J.M., M.I.A.T., H.L., D.R., and R.M.S. conducted experiments and analyzed the data. I.C. and R.C.F. conceived the study and wrote the paper. All authors discussed results and edited the manuscript.

## Author Information

The authors declare no competing financial interests. Correspondence and requests for materials should be addressed to

## Materials and Methods

All procedures were approved under NYU School of Medicine IACUC protocols.

### Behavior: Co-Housing

Pup-naïve C57Bl/6 virgin female mice were bred and raised at NYU School of Medicine and kept isolated from dams and pups until used for these studies when ∼8 weeks old. (For experiments where viral injection was performed, we first allowed two weeks for viral expression before animals were used in experiments.) Dams were initially pre-screened to ensure they behaved maternally, meaning that they retrieved pups and built nests; ∼1% of dams did not retrieve pups and these animals were not used for co-housing. Naïve virgins were initially pre-screened for retrieval or pup mauling before co-housing; ∼5% of the naïve virgins retrieved at least one pup or mauled pups during pre-screening and these animals were excluded from subsequent behavioral studies.

Co-housing of a virgin female with a mother and litter was conducted for 4-6 consecutive days in 80x40x50 cm plastic home cages. The floor was covered with abundant bedding material, food pellets and a pack of hydrogel for hydration placed in a corner of the bin and refreshed daily. Nesting material was also placed in the cage. We first placed the dam and her postnatal day one (P1) litter in the cage. After the dam was acclimated for ∼30 min, we introduced the virgin female with a tail mark for identification. Well-being of the adult mice and pups was monitored at least twice a day. A surveillance infrared camera system (Blackrock Microsystems) was positioned ∼100 cm above the home cage to capture the entire surface. An ultrasonic microphone (Avisoft) was placed in the corner of the cage, ∼10 cm above the nest. (Two initial cages had a second camera placed on the side but these videos were not analyzed for these experiments.)

### Behavior: Pup Retrieval Testing

This test was used for the initial screening of dams and virgin female mice. In addition, outside of the spontaneous home cage behaviors, we specifically monitored pup retrieval every 24 hours by the virgin females (in 2/17 animals co-housed with dam and pups, an additional retrieval test was performed at 12 hrs after co-housing, and neither animal retrieved then). We placed the female mouse to be tested in a behavioral arena (38x30x15 cm) containing bedding and nesting material; the female was alone, without contact with other animals. Each animal was given 20 minutes to acclimate before each testing session began. The entire litter (ranging from three to seven P1-4 pups) were grouped in a corner of the arena and covered with nesting material, and the adult female given an additional two minutes of acclimation (pup group size does not affect retrieval behavior; **fig. S15**). One pup was removed from the nest and placed in an opposite corner of the arena. The experimental female was given two minutes per trial to retrieve the displaced pup and return it back to the nest; if the displaced pup was not retrieved within two minutes, the pup was returned to the nest and the trial was scored as a failure. If the pup was successfully retrieved, the time to retrieval was recorded and the trial was scored as a success. Another pup was then taken out of the nest, placed away from the nest (varying the position of the isolated pup relative to the nest from trial to trial), and the next trial was begun. After ten trials, pups were placed back into their home cage with their dam. We used an ultrasonic microphone (Avisoft) to verify that isolated pups vocalized during testing.

For **Figure 1C**, we used 2-way ANOVAs and Sidak’s multiple comparison tests to compare probability of retrieving in each group over days, and t-test to compare the day of retrieval onset for each group.

### Behavior: Chemogenetic Suppression of Oxytocin Neurons During Co-Housing

*Oxy-IRES-Cre* virgin female mice expressing in OT-PVN cells either <u>D</u>esigner <u>R</u>eceptors <u>E</u>xclusively <u>A</u>ctivated by <u>D</u>esigner <u>D</u>rugs coupled to inhibitory G-protein (DREADDi, also known as hM4D(Gi)) and fused with mCherry, or mCherry alone (as control), were co-housed with mothers and pups as described above. In both experimental (*Oxy-IRES-Cre* virgins expressing DREADDi-mCherry) and control (*Oxy-IRES-Cre* virgins expressing mCherry) cohorts, CNO was administered in the drinking water at a concentration of 25 mg/L. Five sugar pellets were added in each bottle in order to obscure the taste of CNO. At this concentration, for the average mouse body weight (∼30 g) and average daily water consumption in mice (∼6 ml), the average CNO consumption was ∼5 mg/kg per day (http://chemogenetic.blogspot.com/2014/03/cno-in-drinking-water.html).

3/7 animals in the mCherry+CNO cohort had DREADDi virus expressed outside of the PVN due to mis-targeting. These animals were also treated with CNO and as their behavior was similar to the mCherry-expressing animals treated with CNO, we combined data from these groups together in **Figure 3F**.

### Behavior: Video and Audio Analysis

Video and audio recordings were synchronized with the neuronal recordings, and then analyzed with Adobe Audition and Avisoft. For video recordings we used the BORIS suite for scoring of behavioral observations. Two separate teams of independent scorers (two scorers from the Sullivan lab and four scorers from the Froemke lab) were trained in a similar way on how to identify relevant individual and social behaviors during co-housing, and then scored the movies blind to the conditions. The results from each rater were compared and compiled, and results from each lab were cross-validated. Nest entry was considered the moment when the head of the animal entered the nest. Nest exit was considered the time when the rear of the animal left the nest. For **Figure 1D**, we used 1- way ANOVAs and Tukey’s multiple comparison test to compare time in nest across days for each group; we used paired t-tests to compare daily time spent in nest by virgins vs dams.

For determining the distance from nest during shepherding, we measured the distance from the bottom left corner of the cage to the position of the snout of the mouse, and to the position of the nest center. We then calculated distance from the virgin to nest. The start of shepherding was considered to be the moment when the dam makes physical contact with the virgin, and the end of shepherding was the moment when the virgin stops running. In some cases, as the co-housing progressed, we noticed that virgins started running as soon as they noticed the dam approaching; in those cases, the start of shepherding was considered to be the moment when the virgins started running after the dam’s approach. For **Figure 1G**, we used one-sample t-tests to determine if the daily frequency of shepherding was higher than 0. For **Figure 1I**, we used paired t-tests to compare distance from start of shepherding to nest with the distance from end of shepherding to nest.

Audio recordings were processed in Adobe Audition, and isolation/distress calls were distinguished from adult calls and wriggling calls based on the characteristic statistics (bout rate of 4-8 Hz and frequencies of 40-90 kHz).

### Behavior: Observation of Experienced Retrievers

We first confirmed that virgins did not retrieve and dams retrieved at 100% at baseline. The exposures were done in standard behavioral arena (38x30x15 cm). The virgin and dam were acclimated for 20 minutes, then the nest with pups was transferred to this arena. After another 5-10 minutes, we manually isolated one pup at a time so that the dam would retrieve the pup back into the nest. We repeated this 10 times per session. In the experiments where either a transparent or an opaque divided the cage, the two adult animals were acclimated on opposite sides of the barriers. After exposure, the adult animals were separated and the virgins were tested for pup retrieval 30 minutes later as described above. As the preparation for testing and the acclimation to the testing cage also took 30 minutes, this amounted to a total 60 minute interval between virgin observation and testing of responses to isolated pups. The exposure was repeated for four sessions (1/day). A virgin that retrieved at least once during the four days of observation was considered as having acquired pup retrieval behavior. We used chi-square exact tests to compare retrieval between conditions: wild-type mice with no barrier, wild-type mice with transparent barrier, wild-type mice with opaque barrier, and OXTR knockout virgins with transparent barrier.

To determine the role of cortical oxytocin receptor activation during observation of pup retrieval, we implanted cannulas in the left auditory cortex of naïve virgin females (*15*). After animals recovered from surgery, we placed them in the observational chamber, and then infused either saline or the oxytocin receptor antagonist OTA (1 µM solution in saline) through the cannula (**Fig. 5H**). We injected a volume of 1.5 µl at a rate of 1 µl/min. We removed the internal from the cannula five minutes after the end of injection in order to allow the solution to diffuse, and then let the virgins acclimate to the observational chamber for another ten minutes. After this, we exposed virgins to maternal retrievals across the transparent barrier as described above. We used Fisher’s exact test to compare ‘Saline’ and ‘OTA’ conditions (**Fig. 5I**).

### Surgeries: Viral Injections

For testing the effects of DREADDs, stereotaxic viral injections were performed in *Oxy- IRES-Cre* mice (*15,38,39*). Mice were anesthetized with 0.7-2.5% isoflurane (adjusted based on scored reflexes and breathing rate during surgery), placed into a stereotaxic apparatus (Kopf), and bilateral craniotomies performed over PVN (from bregma: 0.72 mm posterior, 0.12 mm lateral). Injections were performed at a depth of 5.0 mm with a 5 μL Hamilton syringe and a 33 gauge needle. Cre-inducible AAV2 hSyn::DIO-hM4D(Gi)- mCherry (University of North Carolina Viral Core) virus was injected into PVN at 0.1 µl/min for a final injection volume of 1.2-1.5 µl. The craniotomy was sealed with a silicone elastomere (World Precision Instruments), the skin sutured. Animals were used for experiments after two weeks to allow for viral expression.

For fiber photometry, we performed viral injections into the left auditory cortex (∼1.7 mm anterior from the occipital suture, 1 mm ventral from skull), using a similar procedure. We injected 1 µl of AAV1 Syn::GCaMP6s (Addgene) at a titer of 1x10^13^ vg/mL in the auditory cortex of wild-type mice. Following the injection, we implanted a 400 µm optical fiber (ThorLabs) just above the auditory cortex, or inserted in the superficial layers of the cortex (200-300 µm below the pial surface).

For imaging PVN→AC projection neurons, we injected 50 nl of AAVrg-hSyn1- GCaMP6s-P2A-nls-dTomato virus (Addgene) at three locations within the left auditory cortex. The virus titer was 1.3xE^12^ vg/ml, and we injected at 10 nl/min.

### Surgeries: Microdrive Implantations

For *in vivo* single-unit electrophysiology we implanted microdrives either in the left PVN (**Figs. 2-4**) or the left auditory cortex (**Fig. 5**). We built microdrives using the parts and instructions for 4-tetrode Versadrives (Neuralynx), adapting these instructions for two bundles each made up of eight 12.5 µm Nichrome wires. The day of the implantation, the wires were cleaned and gold plated to achieve impedances <500 kΩ. After the virgin female mice were anesthetized with isoflurane, a craniotomy (1.5-2 mm in diameter) was performed above the target structure, and two additional small craniotomies were performed in the occipital bone and the right parietal bone for insertion of bone screws. The ground and reference wires of the microdrives were soldered separately to these two bone screws. The dura was removed at the desired implantation site and the electrode bundles were slowly lowered to ∼500 µm above the target brain structure (4 mm ventral from pia for PVN). For recordings from the auditory cortex, we first acutely recorded multiunit activity with tungsten electrodes to localize auditory cortex during implantation procedure. Auditory cortex was found with pure tones (60 dB SPL, 7-79 kHz, 50 msec, 1 msec cosine on/off ramps) delivered in pseudo-random sequence at 0.5-1 Hz. The craniotomy was covered with mineral oil and silicone elastomere, and the microdrive was secured to the skull using dental cement (C&B Metabond).

For optically-identified recordings from OT-PVN units, we used *Oxt-IRES-Cre* mice bred into a C57BL/6 background. Prior to the microdrive implantation, we injected in the PVN of these animals 1-2 µl of AAV1-CAG::DIO-ChR2 at a titer of 1x10^13^ vg/ml. (In one animal we injected AAV2-EEF1::DIO-ChETA at a titer of 1x10^13^ vg/ml.) Then, after implanting and cementing the microdrive, we rotated the head of the animal at a 45°- 50° angle around the anterior-posterior axis, and another craniotomy was done at 0.72 mm posterior and 3.5 mm left from Bregma. A 400 µm optical fiber for delivering blue light was slowly lowered at this position for 4.5 mm below the brain surface. For dual recordings, we implanted a microdrive in left PVN, injected the GCaMP virus (as described above) and implanted an optic fiber in left auditory cortex during the same surgery.

### Surgeries: Optical Fiber Implantation

For fiber photometry we used 400 μm diameter 0.48NA optic fibers inserted into 2.5 mm in diameter ceramic ferrules (Doric Lenses Inc.). For recordings of PVN→AC neurons), we used 1.25 mm metal ferrules, with fibers 1 mm long (below ferrule) for recordings in the auditory cortex and 5 mm long for recordings in PVN. They were implanted during the same surgery as the viral injections, and cemented to the skull as described above.

### Micro-Computed Tomography Imaging

The localization of the implanted electrodes was assessed *in vivo* using micro-computed tomography (μ-CT) scans in post-implanted mice, followed by co-registration with an online digital MRI mouse brain atlas. The µ-CT datasets were acquired using the μ-CT module of a MultiModality hybrid micro-Positron Emission Tomography (µ-PET) /µ-CT Inveon Scanner (Siemens Medical Solutions, Knoxville, TN USA). The Inveon scanner is equipped with a 165-mm x 165-mm X-ray camera and a variable-focus tungsten anode X-ray source operating with a focal spot size of less than 50 μm. The scan consisted of a 20- minute whole-head acquisition over an axial field of view of 22 mm and a transaxial of 88 mm with a resolution of 21.7 μm pixels binned to 43.4 μm. 440 projections were acquired using a 1 mm aluminum filter, a voltage of 80 kV, and a current of 500 μA. The data sets were reconstructed using the Feldkamp algorithm (*40*).

The hybrid scanner is equipped with a M2M Biovet (Cleveland, Ohio USA) module used to monitor continuously vital signs. All mice were monitored continuously throughout the scanning session via a respiration sensor pad (SIMS Graseby Limited, Watford, Hens, UK). The imaging scan consisted of initially placing each mouse in an induction chamber using 3-5% isofluorane exposure during 2-3-min until the onset of anesthesia. The animal was then subsequently positioned laterally along the bed palate over a thermistor heating pad in which 1.0% to 1.5% isofluorane was administered via a 90° angled nose cone throughout the scan. The head of each subject was judiciously oriented perpendicular to the axis of the mouse body so that the extracranial part of the implanted electrode could be easily kept away from the field of view of the µ-CT image acquisition. Importantly, the large extracranial metal components and dental cement of the implant can cause beam hardening that can appear as cupping, streaks, dark bands or flare in the µ-CT (*41–43*). To this effect, the head positioning helped reduce the risks of image artifacts that could be induced by the implant along the path of the X-ray beam.

Unlike MRI, μ-CT imaging can be performed on subjects with metal implants. However, lack of the soft tissue contrast of the µCT limited its usefulness to provide the needed brain anatomical detail in order to verify the electrode correct localization. Our approach combined registration of post-implant micro-computed tomography (µ-CT) with an existing online MRI brain atlas for adult C57Bl/6 mice from the Mouse Imaging Centre (https://wiki.mouseimaging.ca/display/MICePub/Mouse+Brain+Atlases).

The three-dimensional MRI mouse brain atlas was established by acquiring 40 individual *ex vivo* mice using T2-weighted sequence on a 7-Tesla scanner. All the data were averaged and resulted into a 40 µm isotropic resolution dataset detailed in Dorr et al. (*44*). The hypothalamic PVN was manually segmented and color-coded, with the guidance of the P56 coronal Allen mouse histology brain atlas (*45*). This region was set as the target of reference. A rigid co-registration between µ-CT and the modified MRI atlas images was systematically performed using a commercial software Amira (Thermo Fisher Scientific, Waltham, MA USA). Both datasets were then overlaid for visual analysis and to determine the sub-millimetric localization of the electrode tip.

### Single-Unit Recordings During Behavior

Neuronal recordings were performed using the Cereplex µ headstage, a digital hub and a neuronal signal processor (Blackrock Microsystems). The recordings were synchronized with a video recording system (Neuromotive, Blackrock). The optical fiber was connected to a blue laser triggered using an analog output from the recording system. The laser pulses were 5 msec in duration and delivered at either 2 or 5 Hz. The light intensity was controlled by adjusting the output of the laser, and at least three different intensities were tested in each animal. To allow free movement of the implanted mice, we used a pulley combined with a fiber-optic and electrical rotary joint/commutator (Doric Lenses). Before the start of the experiment the electrode bundles were lowered into the target structure and then daily advanced by ∼70 µm. For PVN recordings, optical tagging tests were performed daily.

Neurons were isolated using the Cerebrus and Boss software (Blackrock Microsystems). For identifying oxytocin neurons in optical tagged recordings we aligned neuronal activity to the onset of the blue light pulse. Neurons that reliably fired after the onset of blue light were selected as being oxytocin neurons. For analyzing changes in firing patterns, we aligned the spike trains to the onset of the behavioral episode, and calculated the firing rate for the duration of the behavior. In the case of ‘nest entry’, we used a 40 second cutoff, in order to avoid contamination of the data with modulations that might be caused by other behaviors in the nest (sleeping, nest building, etc). For each behavioral episode we also calculated the corresponding baseline firing rate, either for an interval similar in duration with the behavioral episode (in the case of nest entry), or for 10 seconds prior to the start of the behavioral episode (‘for shepherding’ and ‘dam retrievals’). The change in firing rate was calculated as: (Spiking behavior – Spiking baseline)*100/(Spiking behavior + Spiking baseline) (**Fig. 2E,F,I,J**; **Fig. 3C**). To determine if these changes in firing rate were statistically significant, we shifted the behavior intervals by a random value between -500 and 500 seconds, and then we calculated new firing rates and modulation percentages for 1000 random shifts. All cells for which the modulation of activity in the unshifted data was greater than modulation in 950 shuffles were considered to contain a significant peak of activity (*46*). A similar analysis was done for PVN spike data recorded both during the observation of maternal retrieval and during the execution of the retrieval by the virgin, in the dual implanted animals (**Fig. 5D,E**). To investigate how PVN activity correlates with activity in the auditory cortex, a population average change in PVN firing was calculated for each maternal retrieval trial of the observational learning procedure.

For quantifying unit activity during behavioral episodes, we aligned the neuronal data to the onset of the behavior, and binned the spiking data in 250 msec bins for an interval between -20 seconds and +40 seconds (-10 to +20 in the case of shepherding episodes). We then calculated the z-score value for each bin in the chosen time window; for display purposes, we used bounds of -3 (min) and +3 (max). For single-unit recordings in the auditory cortex, the spike trains were aligned to the onset of the played-back pup call. The average firing rate during the pup call (1 second interval) was normalized to the firing rate during baseline (1 second preceding pup call onset).

### Fiber Photometry

To perform fiber photometry, we connected the optic fiber implanted in the auditory cortex and housed in a ceramic ferrule to a custom-built photometry rig (*47*). A 400 Hz sinusoidal blue light was delivered via the optical fiber from an LED (30 µW) for GCaMP6s excitation. We collected the emitted (green) light via the same optical fiber, used a dichroic mirror and appropriate filters to direct emitted light to a femtowatt silicon photoreceiver (Newport), and recorded using a real-time processor (RX8, TDT). The analog readout was then low-pass filtered at 20 Hz. The intensity of blue light excitation was adjusted to produce similar baseline fluorescence levels across sessions in the same mouse. The sound processor used for delivering the pup calls (RZ6, TDT) was synchronized with the fiber photometry system.

For investigating changes in neuronal activity of the left auditory cortex throughout co-housing, we recorded responses to six different played-back pup calls every 24 hours. To avoid possible changes in the magnitude of calcium transients with changes in the position of the mouse relative to the speaker, in some recordings we restrained the mouse in a mash cup, at the same distance from the speaker. Data from restrained or unrestrained mice showed similar trends, and were thus pooled together. To determine the evoked response recorded with fiber photometry, we calculated the ΔF/F for each pup call interval.

### Anatomy

To verify viral expression, at the end of the experiments, animals were fixed with transcardiac perfusions of 4% paraformaldehyde. Brains were removed and further preserved in a paraformaldehyde solution for 1-2 hours at 4° C. Afterwards, brains were sequentially cryopreserved in 15% and then 30% sucrose solution, embedded in OCT solution and sectioned (30 µm) with a cryostat then mounted on positively-charged slides. Immunohistochemistry was performed on the mounted sections as previously described (*48, 49*). Sections were blocked in 5% goat serum solution for an hour at room temperature or for 14 hours at 4° C. A solution of the appropriate primary antibodies was diluted in 1% goat serum and 0.01% Triton X solution and then applied for 24 hours at 4° C. We used a rabbit anti-oxytocin antibody (EMD Millipore, 1:500), mouse anti-oxytocin antibody (a gift from Dr. Harold Gainer at NIH), a chicken anti-GFP (Aves, 1:500) and a chicken anti-mCherry (Abcam, 1:1000). The sections were washed in PBS solution, and a solution of fluorophore-conjugated secondary antibodies applied for 1.5 hours at room temperature. All secondary antibodies were from Jacksons Immunoresearch and used 1:200. Slides were examined and imaged using a Carl Zeiss LSM 700 confocal microscope with four solid-state lasers (405/444, 488, 555, 639 nm) and appropriate filter sets. For imaging sections co-stained with multiple antibodies, we used short-pass 555 nm (Alexa Fluor 488), short-pass 640 nm (Alexa Fluor 555), and long-pass 640 nm (Alexa Fluor 647) photomultiplier tubes.

---

**Fig. S1.**
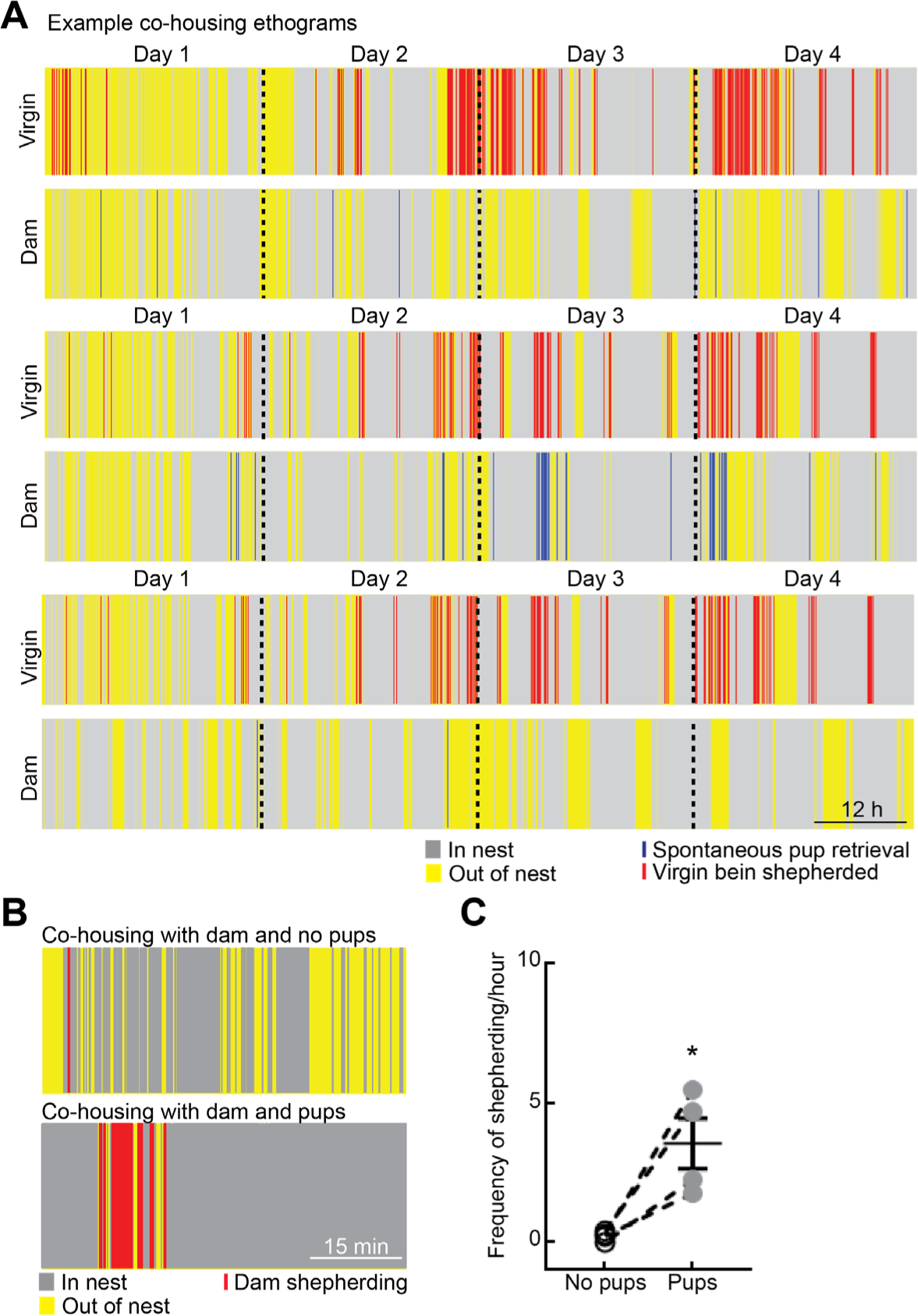
Dynamics of voluntary nest entry and shepherding during co-housing. (A) Additional ethograms of time in or out of nest for three other dam-virgin dyads. (B) Shepherding behavior requires pups in nest. Example ethograms from two separate virgins, co-housed without pups (top) or with pups (bottom). Note difference between time in nest and amount of shepherding when pups were present. (C) Summary of shepherding events per hour when pups were removed vs when pups were present (‘No pups’: 0.2±0.1 events/hour; ‘Pups’: 3.5±0.9 events/hour, N=4, p=0.03, Student’s two-tailed paired t-test).

**Fig. S2.**
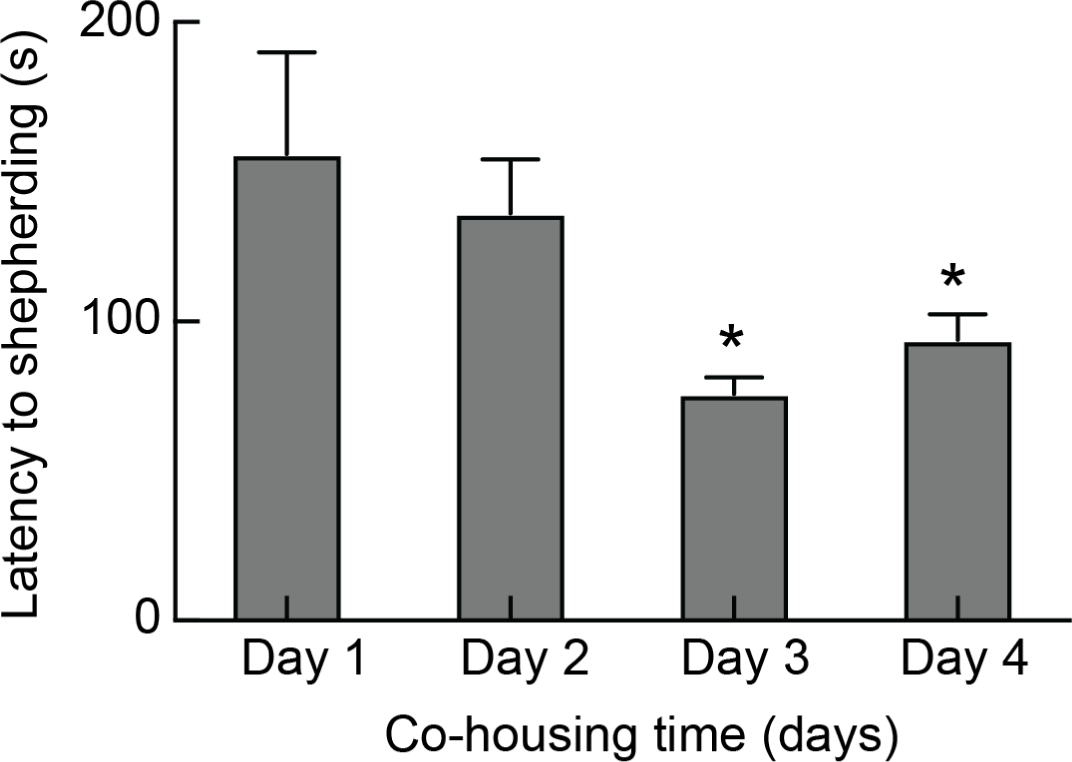
Average latency for dams to shepherd virgin to the nest (time 0 is virgin nest exit). Latency decreased over days of co-housing (day 1: 155.9±34 s; day 2: 136.1±17.94 s; day 3: 75.7±5.5 s; day 4: 93.9±8.5 s; n=676 events from N=9 dam-virgin pairs, p<0.0001, one-way ANOVA, Holm-Sidak’s multiple comparison test).

**Fig. S3.**
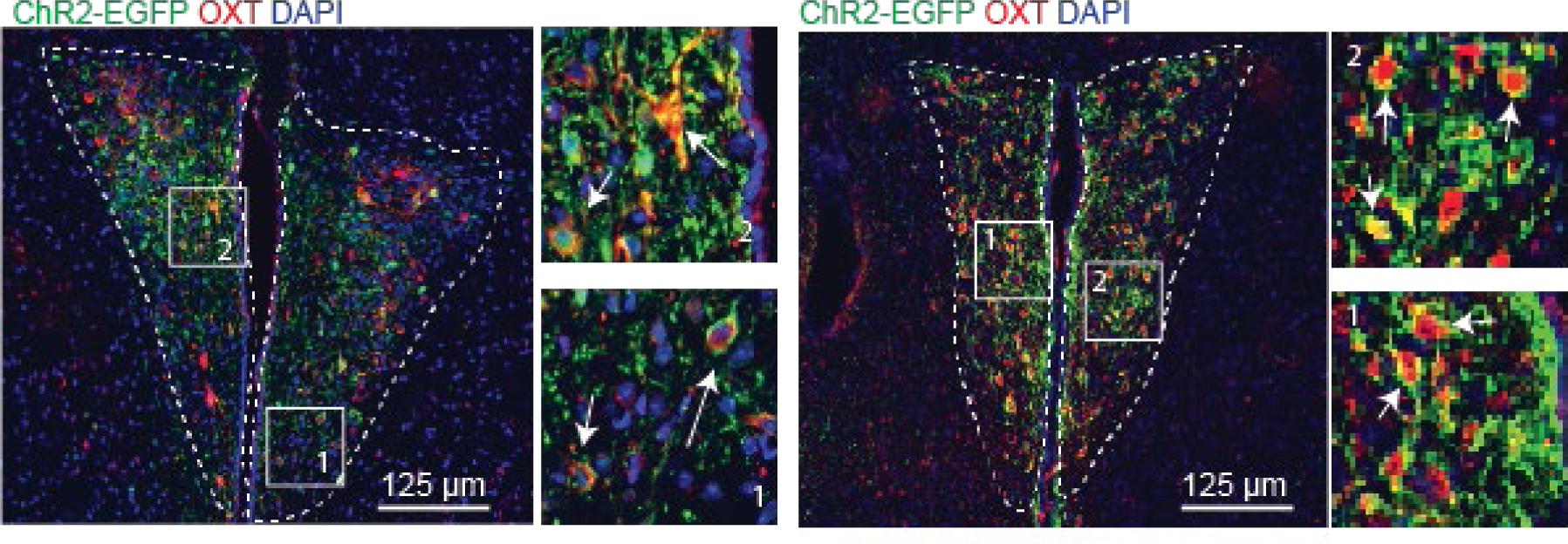
Example immunostained tissue sections of PVN showing co-labeling of oxytocin peptide (red) and ChETA channelrhodopsin-2 conjugated to EGFP (green) together with DAPI labeling for cell density (blue). Insets show 2x magnified images to highlight the co- localization of red and green staining (arrows) in PVN cells. 84.8±7.2% of cells (N=6 animals) expressing ChETA also expressed oxytocin peptide.

**Fig. S4.**
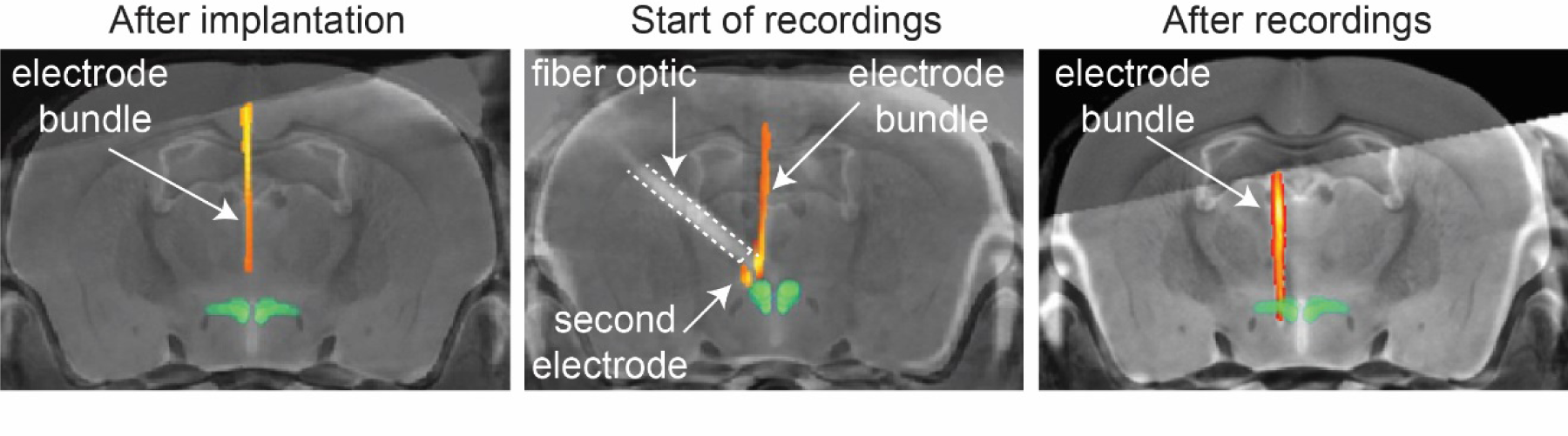
Verification of electrode locations in PVN with combined CT/MRI. CT imaging was performed after implantation and subsequently co-registered with a mouse MRI atlas to localize electrode bundles within PVN, as well as the separate fiber optic implanted for optogenetic identification of oxytocin neurons. Electrodes were then lowered at the end of each day of co-housing. Yellow/orange, CT imaging of electrodes in situ. Images shown are from different animals.

**Fig. S5.**
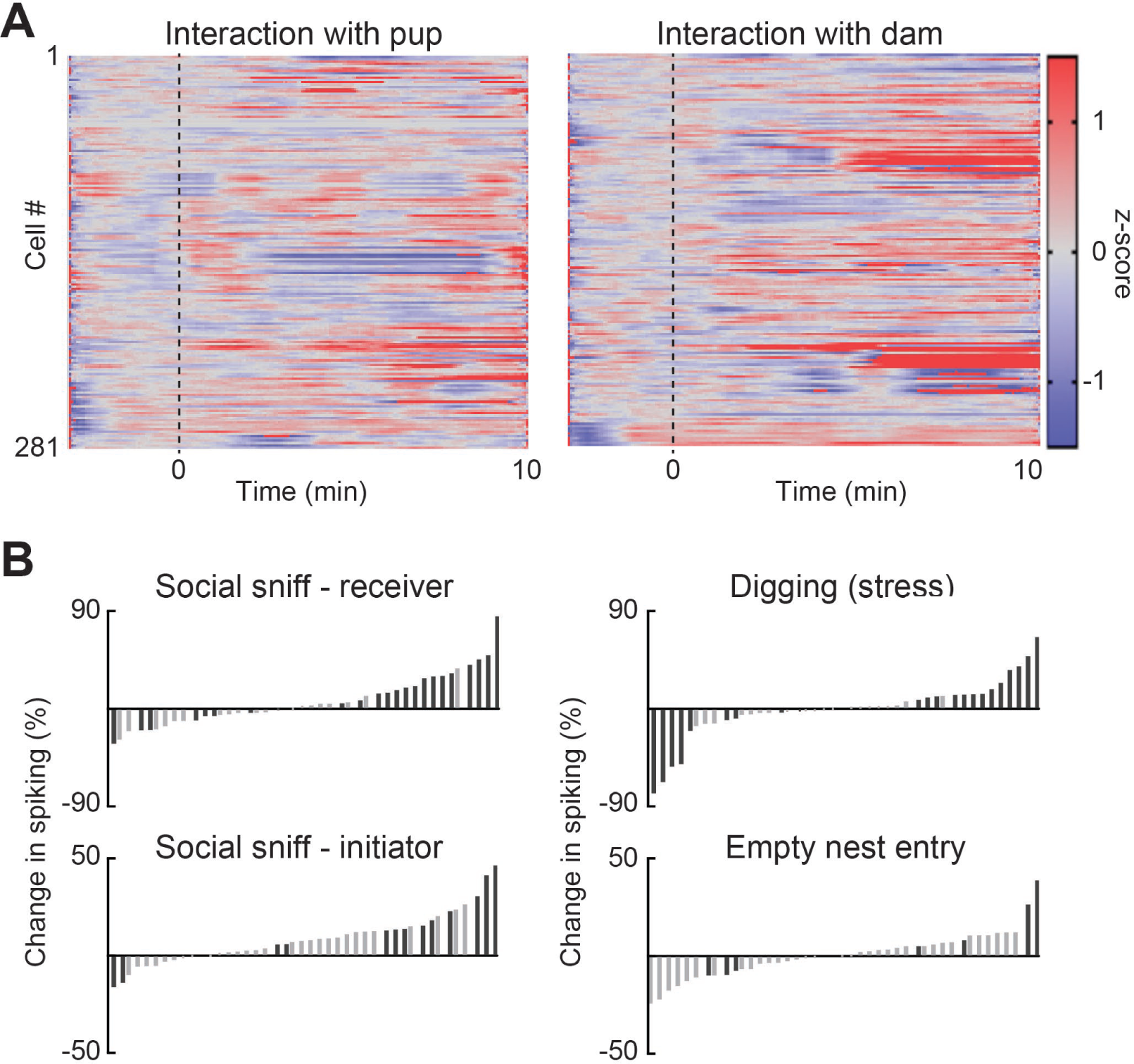
PVN neurons in adult female virgin mice respond more robustly during short-term interactions with dams than with pups. (A) Normalized (z-scored) spiking for 281 PVN neurons recorded during short interactions with pup (left) or a dam (right). Recordings were aligned to start of interaction. (B) Social sniffing between the virgin and dam activate PVN neurons (15/43 neurons significantly activated while the virgin was sniffed; 11/43 neurons activated while the virgin sniffed the other adult). A similar fraction of PVN neurons was activated (13/43) during digging. Importantly, only a few PVN neurons were activated by entering an empty nest without pups (4/43). Black bars, significantly modulated neurons.

**Fig. S6.**
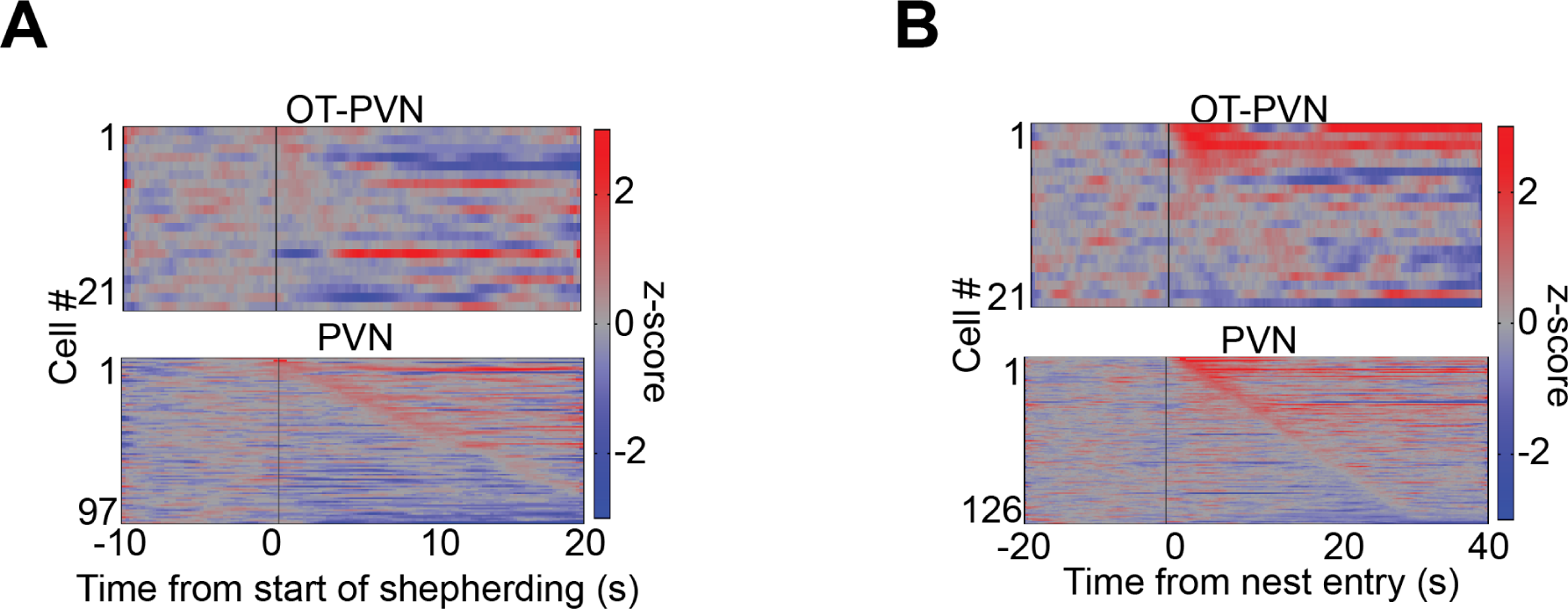
Shepherding and nest entry activate identified OT and other PVN cells. (A) Z-scored firing rates of OT-PVN and PVN cells during shepherding. (B) Z-scored firing rates of OT- PVN/PVN cells during nest entry.

**Fig. S7.**
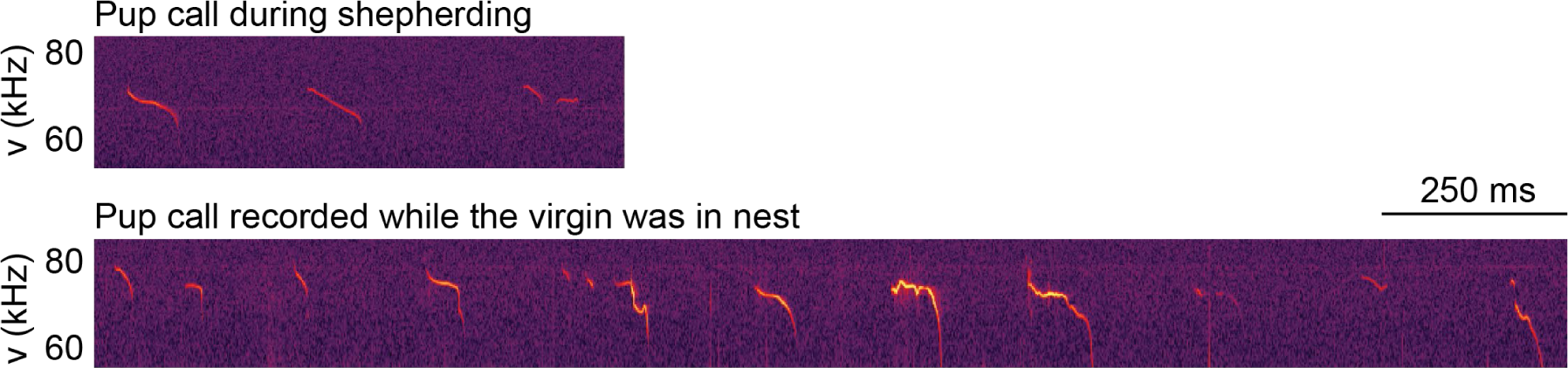
Pups vocalize and make distress calls in home cage during shepherding and nest entry. Shown are example spectrograms over time.

**Fig. S8.**
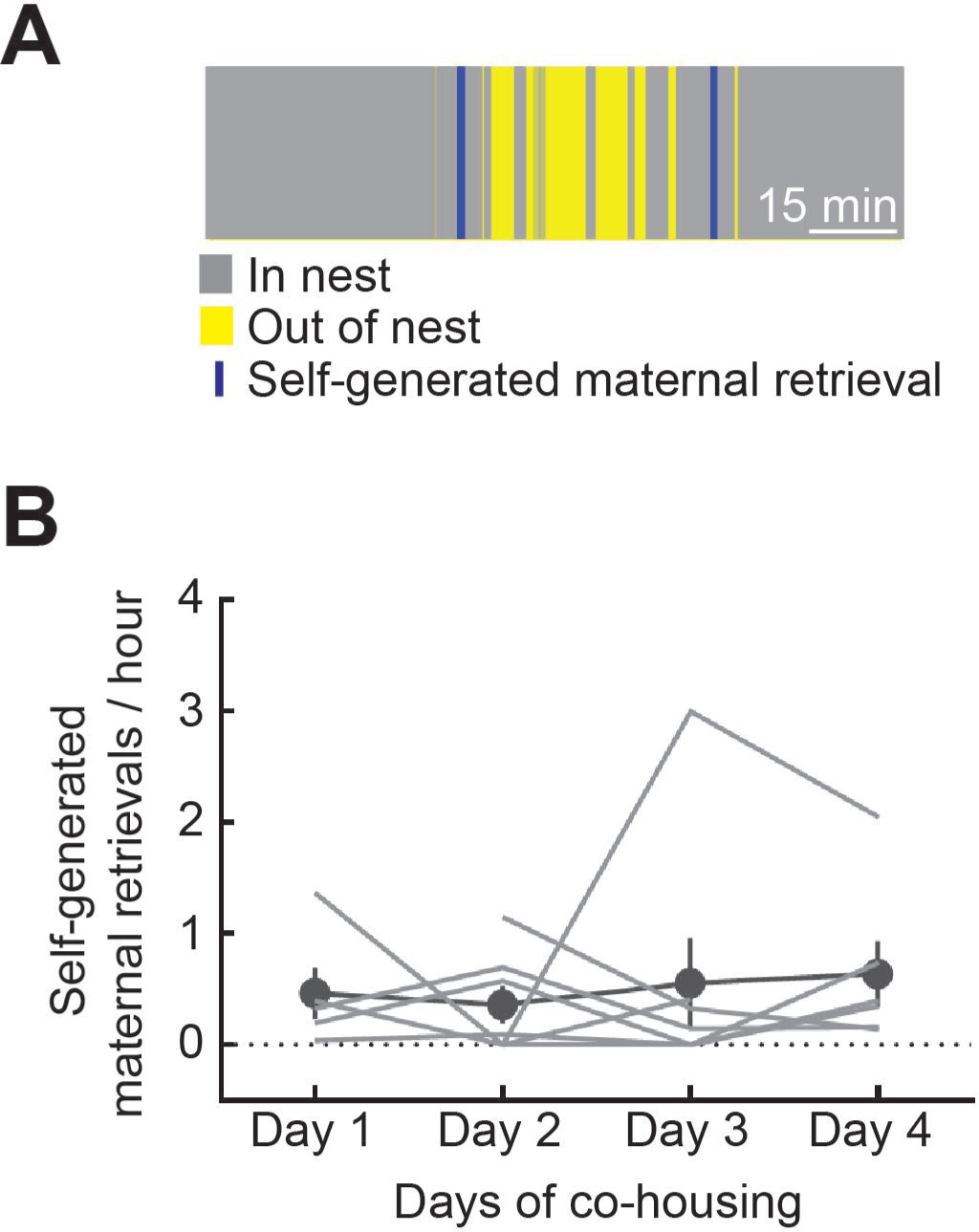
Frequency of self-generated retrievals by dams. (A) Example ethogram depicting two separate spontaneous retrievals by the dam. (B) Summary of individual dam spontaneous hourly pup retrieval rate per day (day one: 0.5±0.2 events/hour, day two: 0.4±0.2 events/hour, day three: 0.6±0.4 events/hour, day four: 0.6±0.3 events/hour, N=7). Lines, individual dam behavior. Filled circles, daily average across animals.

**Fig. S9.**
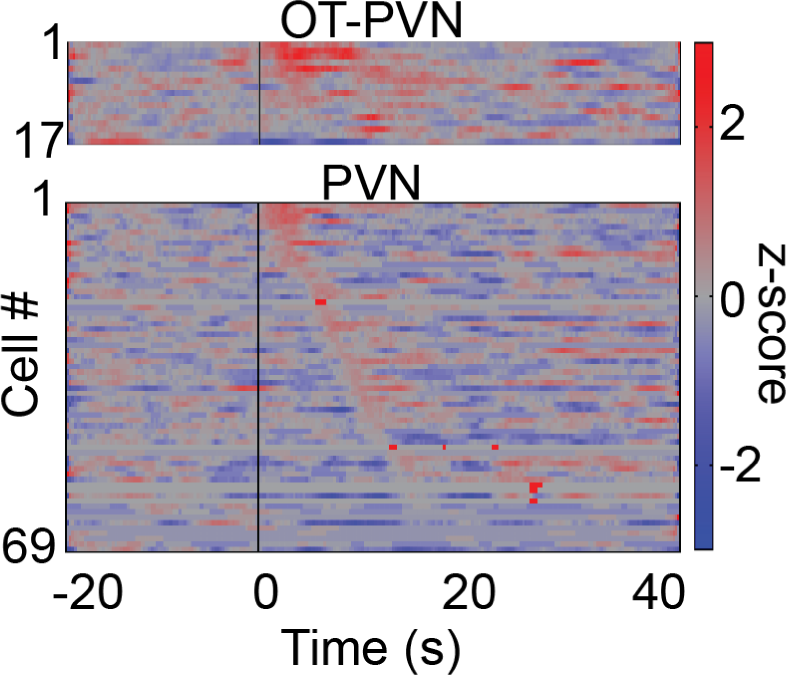
Maternal retrievals activate identified OT and other PVN cells in virgins. Z-scored firing rates of virgin OT-PVN and PVN cells recorded during spontaneous retrieval episodes by dams. Time 0, start of maternal retrieval.

**Fig. S10.**
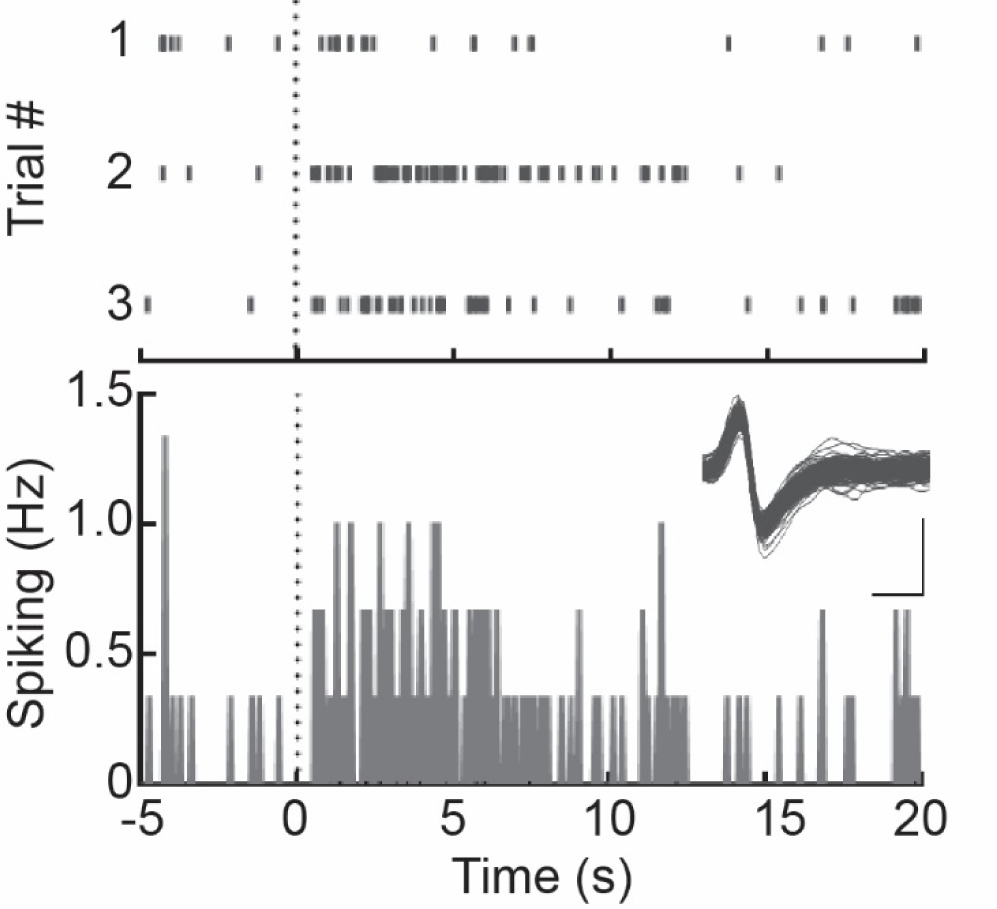
Virgin PVN responses when mother brought pups to virgin. For these animals, three separate pup delivery episodes occurred in a twenty-minute period after the second day of co-housing. On each of those occurrences, this PVN unit recorded in the virgin responded with a burst of spikes for several seconds until the dam returned to retrieve the pup herself. Top, raster of each of these three trials. Bottom, summary PSTH. Inset, spike waveforms; scale bar: 0.4 ms, 0.5 mV.

**Fig. S11.**
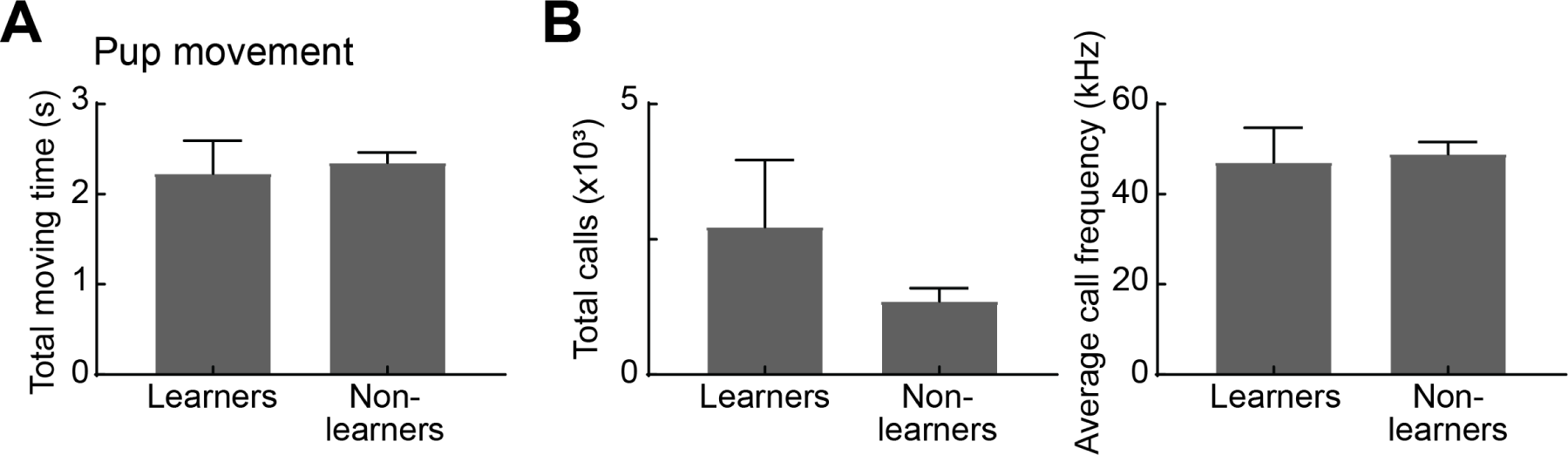
No significant differences in behavior of pups between virgins that began retrieving after observation (‘Learners’) and those virgins that did not begin retrieving after observation (‘Non-learners’). (A) Pup movement during observational learning is not significantly different between the two groups. (B) Pup vocalizations during observational learning do not differ in call rate and frequency between groups.

**Fig. S12.**
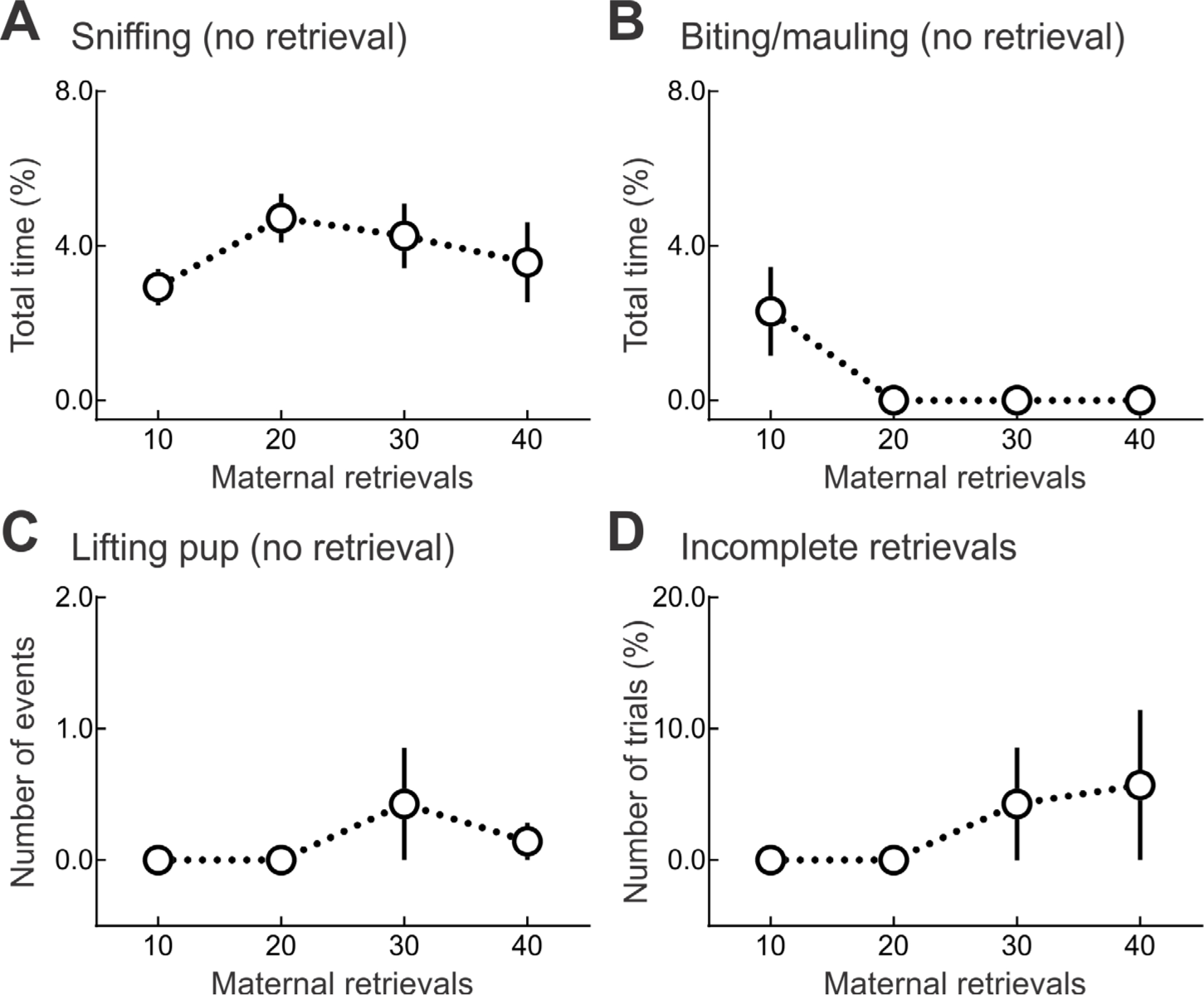
Pup retrieval errors by inexperienced virgins, measured after each observation session of 10 retrievals by dam each day. (A) Average time spent sniffing pups by virgins during retrieval testing was similar across days (day one: 2.9±0.5% total time spent sniffing, day two: 4.7±0.6% total time, day three: 4.3±0.8% total time, day four: 3.6±1.0% total time, N=7). (B) Biting or mauling pups was only observed on the first day (day one: 2.3±1.2% total time spent biting/mauling, days two to four: 0.0±0.0% total time, N=7). (C) Lifting pup and putting it back in place was observed on days three and four (days one and two: 0.0±0.0 observed episodes of pup lifting, day three: 0.4±0.4 observed events/animal/session, day four: 0.1±0.1% observed events/animal/session, N=7). (D) Incomplete retrieval was defined as relocating the pup somewhere other than the nest, and was observed only on days three and four (days one and two: 0.0±0.0% incomplete trials, day three: 4.3±4.3% trials incomplete, day four: 5.7±5.7% trials incomplete, N=7).

**Fig. S13.**
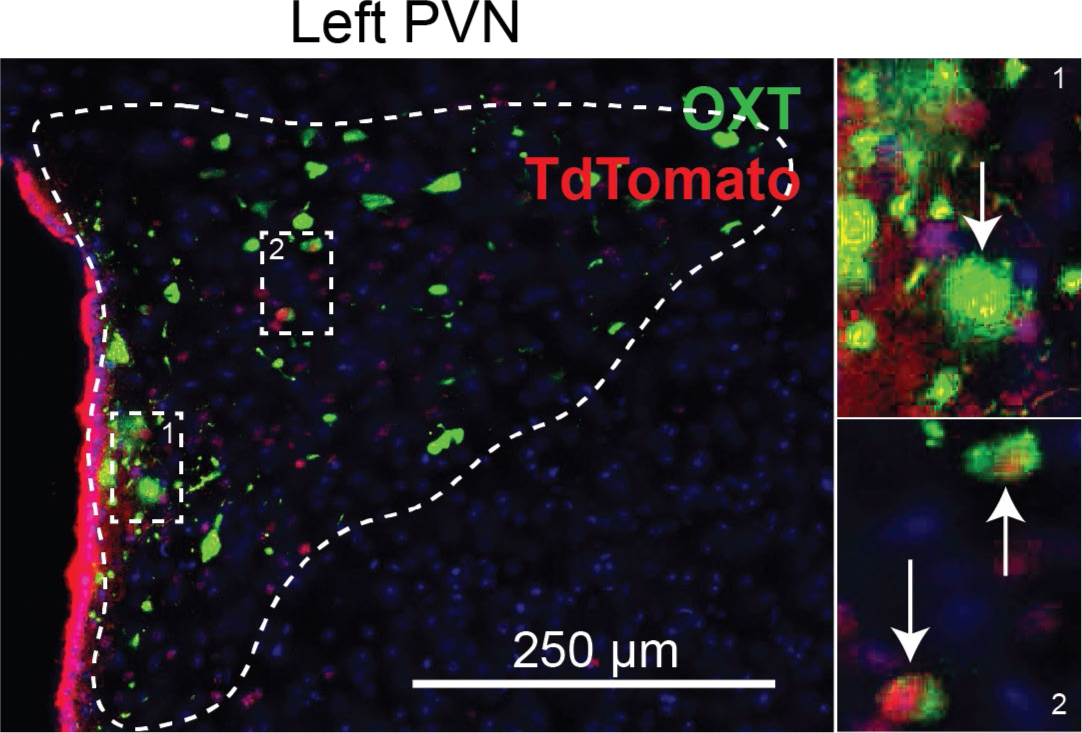
Example immunostained tissue section of PVN showing co-labeling of oxytocin peptide (green) and GCaMP6s expressed together with TdTomato (red).

**Fig. S14.**
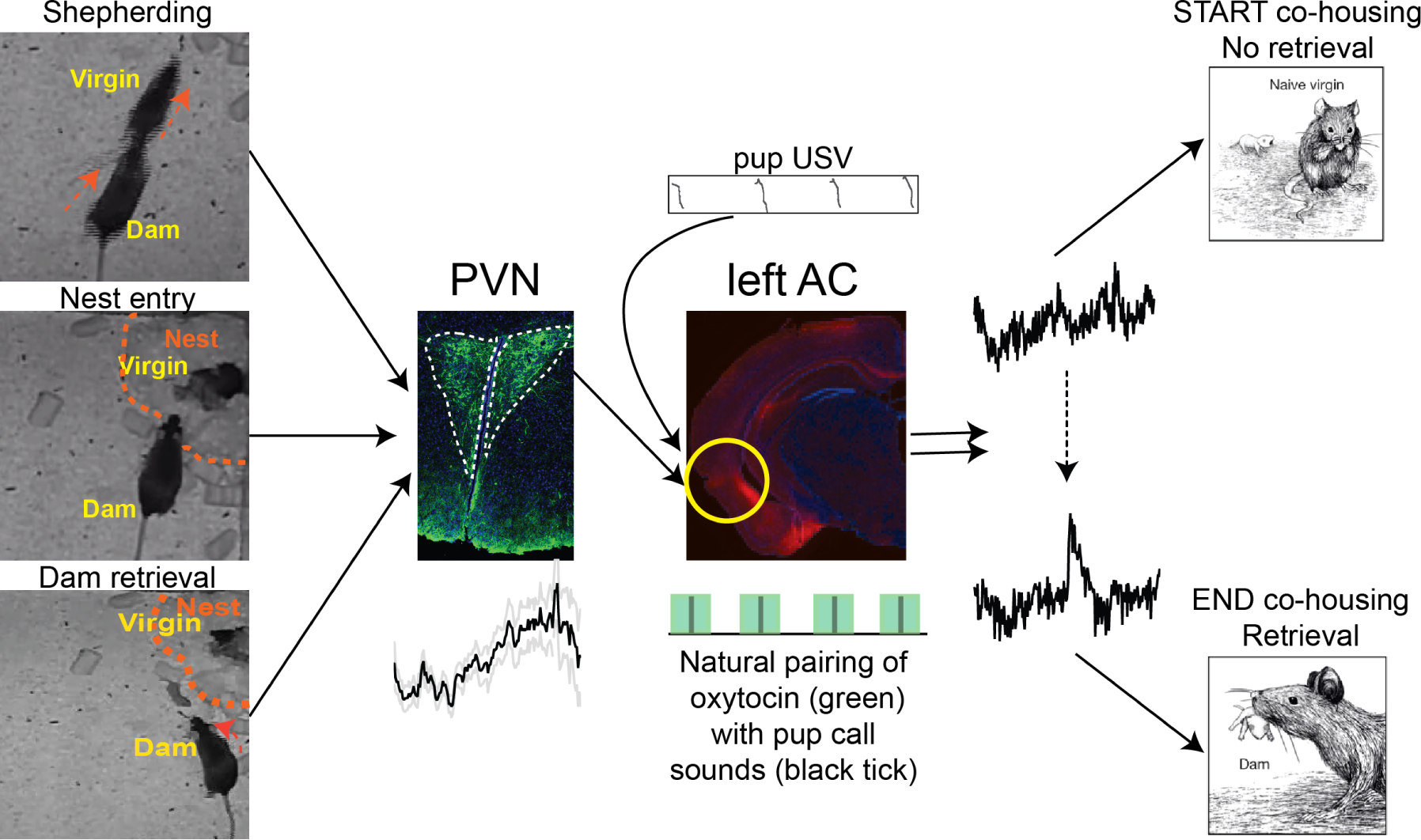
Schematic of overall hypothesis connecting behavioral interactions between dam and virgin to activation of PVN and OT-PVN neurons and subsequent modulation/modification of left auditory cortex for reliable recognition of pup distress calls.

**Fig. S15.**
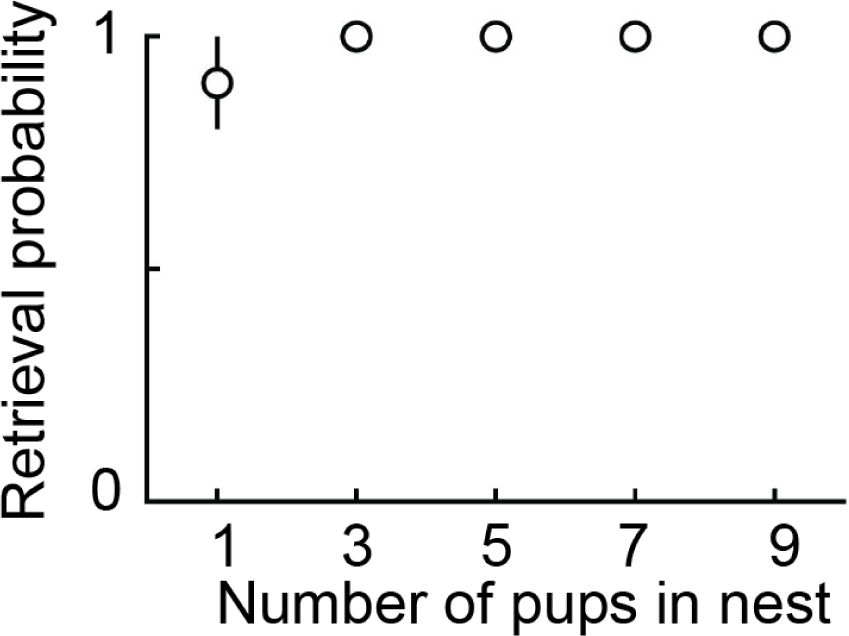
Pup retrieval by experienced animals (N=3) is independent of litter size (i.e., number of pups in nest).

## Supplementary Movies

Movie S1

Example of spontaneous pup retrieval by mother occurring during co-housing (same as in Fig. 3A).

Movie S2

Example of virgin spending time in nest with pups after days of co-housing; same dam- virgin dyad recorded on days one through four.

Movie S3

Retrieval onset in virgin co-housed with dam and pups. Example pup retrieval testing on day 0, day 1, and day 3.

Movie S4

Retrieval onset in virgin co-housed with pups. Example pup retrieval testing on day 0, day 1, and day 5.

Movie S5

Example of dam shepherding virgin back to nest (same event as in **Fig. 1E**).

Movie S6

Movie with more examples of dam shepherding from different co-housing pairs.

Movie S7

Movie with more examples of self-generated retrievals from different co-housing pairs.

Movie S8

Example of dam bringing pup near virgin.

Movie S9

Testing social transmission of maternal behavior with transparent barrier separating virgin from dam and pups; after two days, virgin was retrieving.

Movie S10

Testing social transmission of maternal behavior with opaque barrier separating virgin from dam and pups; virgin did not begin retrieving even after four days.

Movie S11

Pup retrieval errors by virgins before learning to correctly retrieve.

## Notes

#### Summary of Updates

Response to first round of reviewer comments.

